# Nar1 binds the cytosolic iron sulfur cluster assembly targeting complex via a bipartite interaction interface

**DOI:** 10.64898/2026.03.06.710161

**Authors:** Jackson V. Ho, Anastasiya Buzuk, Melissa D. Marquez, Beatrice Wang, Deborah L. Perlstein

## Abstract

The cytosolic iron-sulfur cluster assembly (CIA) pathway maturates essential nuclear and cytosolic Fe–S proteins required for genome maintenance and cellular metabolism. Nar1 (also called CIAO3 or IOP1) is a conserved Fe–S protein that connects the early and late steps of the CIA pathway, yet the molecular basis for its proposed function as a metallocluster carrier remains poorly defined. In particular, the interactions responsible for Nar1 recruitment to the CIA targeting complex (CTC) during cluster delivery remain unknown. Here, we define the molecular basis for Nar1 recruitment to the CTC using biochemical reconstitution, quantitative protein-protein interaction assays, and AlphaFold modeling. Our data reveal that Nar1 binds the CTC through two distinct interfaces. A primary interface comprises an electrostatic interaction that anchors Nar1 to a conserved acidic surface on the Cia1 subunit of the CTC and a secondary interface involves binding of Nar1’s divergent targeting complex recognition peptide at the Cia1-Cia2 interface. Thus, Nar1 engages a conserved CTC surface that serves as a recruitment platform for multiple binding partners, including CIA clients. Computational structural models position the putative Fe–S cluster donor site of Nar1 adjacent to a proposed acceptor site on Cia2, suggesting that this bipartite binding mechanism positions Nar1 for transfer of an Fe–S cluster to the targeting complex. Together, these findings resolve conflicting models for Nar1 recruitment and establish a mechanistic framework for understanding how the CTC engages multiple binding partners during cytosolic iron-sulfur protein maturation.

## INTRODUCTION

Iron-Sulfur (Fe–S) proteins participate in a wide variety of fundamental cellular processes across all domains of life.^1-4^ Due to the fragility of these cofactors in aerobic environments and the toxicity and scarcity of free iron and sulfide in vivo, the maturation of Fe–S enzymes requires a complex network of conserved and essential metallocofactor biogenesis factors that assemble, traffic, and target nascent Fe–S cofactors, facilitating maturation of apo-client proteins. In eukaryotes, Fe–S cluster biogenesis occurs across multiple subcellular compartments, operating in the mitochondria, in cytosol and nucleus, and, when present, plastids.^1-3^ The cytosolic iron-sulfur cluster assembly (CIA) system maturates cytosolic and nuclear Fe–S proteins, including primase, DNA and RNA polymerases, helicases and glycosylases. Consequently, inhibition of CIA leads to genome instability, defects in telomere maintenance, increased mutation rates and replicative stress, and contributes to diseases including neuromuscular degeneration and cancer.^5-8^

In the late stages of the CIA pathway, Fe–S clusters are delivered to clients via the CIA targeting complex (CTC; **Figure 1A**).^1,9-11^ This multiprotein complex is proposed to act as the central delivery hub, accepting a nascent cluster from the upstream trafficking factor Nar1 (also known as CIAO3, IOP1 or NARFL in humans) and subsequently delivering the metallocluster to a CIA client.^12,13^ Structural and biochemical studies have begun to reveal how the CTC subunits Cia1 (CIAO1), Cia2 (CIAO2B/FAM96B/MIP18, herein called Cia2b), and Met18 (MMS19) assemble to form the CTC and recruit CIA clients.^9,11,14-16^ Recent structural and biochemical studies have indicated two primary points of contact between the CTC and clients, one between a conserved surface along the side of Cia1’s third β-propeller domain and a second located in the N-terminal domain of Met18.^16-18^ Additionally, many CIA clients contain a conserved targeting complex recognition (TCR) peptide, which binds a pocket formed at the conserved interface between Cia1 and Cia2 to recruit the CTC via binding at the interface of its Cia1 and Cia2 subunits (**Figure 1A**).^19-21^ Despite these advances, comparatively little is known about the interactions that govern recruitment of Nar1, the putative CIA trafficking factor acting upstream of the CTC.^12,22-25^

**Figure 1.**
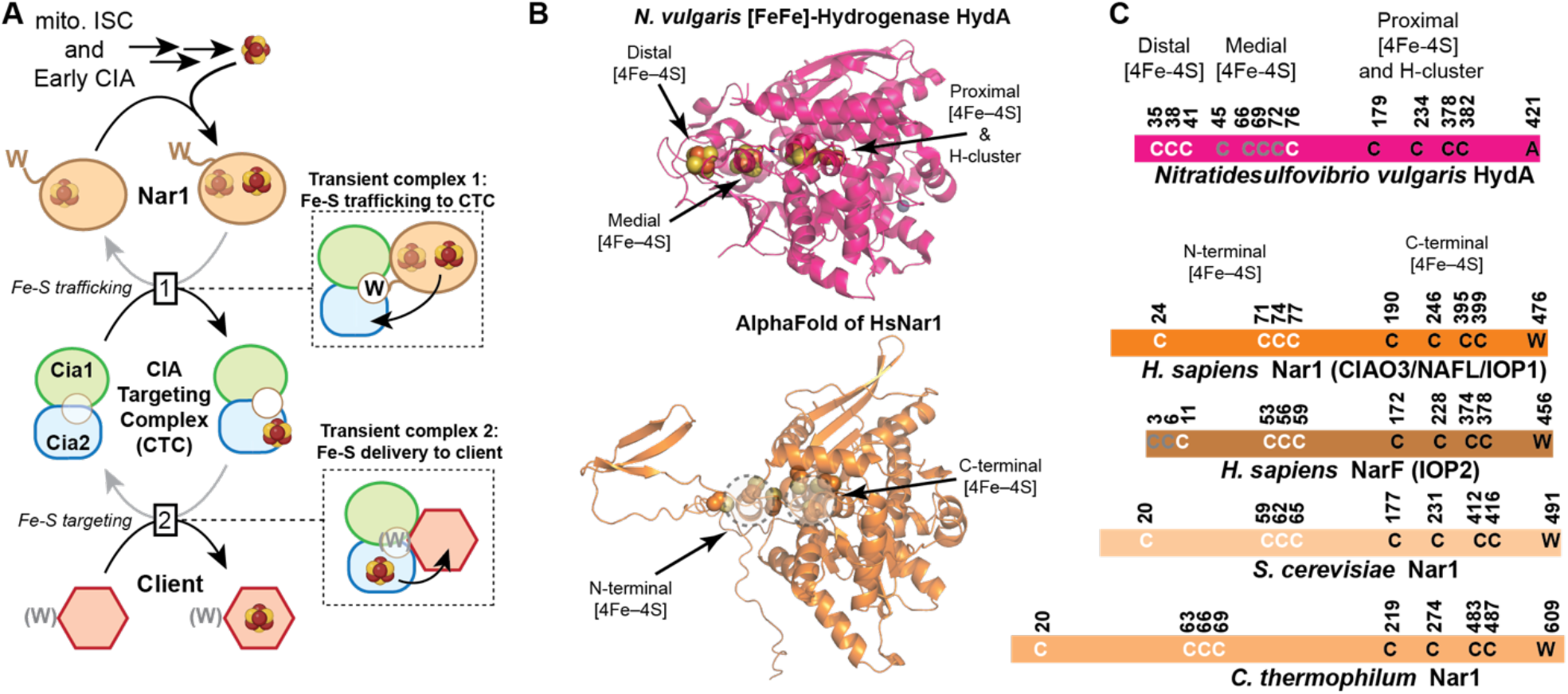
Overview of the CIA pathway and the structural features of Nar1. **A**) Schematic of Fe-S cluster transfer in the CIA pathway. The mitochondrial ISC system and early CIA factors synthesize a [4Fe-4S] cluster, which is transferred to Nar1. Next, Nar1 delivers the cluster to the CTC through formation of a transient complex, and interaction facilitated by Nar1’s C-terminal tail ending in a tryptophan (W) residue. The [Fe-S]-bound CTC engages apo-clients, some of which end in a TCR peptide, to complete the cluster maturation pathway. The MMS19/Met18 subunit of the targeting complex is omitted for clarity. **B**) Structural comparisons of [FeFe]-hydrogenase^26-28^ and *Hs*Nar1, AlphaFold prediction.^29^ Proposed [Fe-S]-ligating cysteine residues of Nar1 are spheres. Dotted circles indicate approximate positions of Nar1’s N-terminal and C-terminal Fe-S binding sites. **C)** Conserved cysteines of hydrogenase are shown, colored white, grey or black for the distal, medial and proximal [4Fe-4S] clusters shown. Nar1 homologs from *Hs* and *Sc* are shown, with their relative positioning and conservation with hydrogenases indicated by the position. The cysteines proposed to ligate the N-terminal and C-terminal Fe-S cluster of Nar1 are colored white and black, respectively. NarF sequences, found in certain vertebrates, have two additional conserved cysteine residues at their N-terminus, colored grey.

Efforts to delineate the Nar1-CTC interaction interfaces have yielded both organism-specific findings and mechanistic discrepancies. Earlier biochemical studies with the human system suggested that Nar1 interacts primarily with the Cia1 subunit of the targeting complex, whereas later studies indicated that the Cia2 subunit was required for Nar1 to associate with Cia1.^19,25,30^ A clue about potential interaction interfaces could be gleaned from the fact that Nar1 terminates in a conserved tryptophan that is required for Nar1’s essential role in the CIA pathway in vivo, suggesting that, like some CIA clients, Nar1 also possesses a targeting complex recognition (TCR) peptide.^20^ Later in vitro studies showed that Nar1 competes with the CIA client Leu1, which terminates in a TCR motif, for binding to the CTC, further corroborating the hypothesis that the CTC functions via a ping-pong mechanism, where Nar1 binds and delivers a nascent cluster to the CTC, followed by formation of a CTC-client interaction and maturation (Figure 1A).^19^ However, previous immunoprecipitation-mass spectrometry studies suggested that the CTC could be bypassed, allowing Nar1 to bind CIA clients directly and presumably deliver an Fe-S cluster to them. Still other studies have suggested that Nar1 can bind to and receives its Fe-S cluster from the CIA scaffold in the early stage of the pathway whereas others have suggested the mitochondrial ISC trafficking protein ISCA1 can also be an on-pathway donor for Nar1’s cluster.^31^ Further complicating the picture, Rouault and coworkers have proposed that an ISCU/HSC20 complex, not Nar1, provides Fe-S clusters to the CTC.^32^

Nar1 itself is particularly intriguing because of its evolutionary relationship to [FeFe]-hydrogenases, ancient enzymes involved in hydrogen metabolism, which contain multiple Fe–S clusters that relay electrons to and from the catalytic H-cluster (**Figure 1B-C**).^33-35^ Nar1 retains eight conserved cysteine residues positioned analogously to residues that ligate the proximal and medial [4Fe–4S] clusters of hydrogenase.^22,36^ These residues are therefore proposed to coordinate two [4Fe– 4S] clusters within Nar1, and possibly a third [2Fe–2S] cluster bound in place of the H-cluster.^36^ Unlike the stably-bound Fe–S clusters of hydrogenase that function in electron transfer, at least one or more of the Nar1 metalloclusters are proposed to be transiently associated, allowing Nar1 to serve as an intermediate carrier proposed to link the early-acting CIA scaffold with the late-acting CTC. Structural insight into Nar1 has been limited, in part because isolation of fully cluster-loaded, homogeneous protein has proven difficult, leaving the molecular architecture of Nar1 and its interactions with the targeting complex poorly defined. When combined with the lability and oxygen sensitivity of these clusters, efforts to determine the number, nuclearity and functional roles of these Fe–S cluster cofactors have been challenging to decipher. Additionally, some animals encode a second Nar1-like paralog, called NARF or IOP2, the role of which remains largely unknown (**Figure 1C**).^13,34^ The many contradictory findings across prior studies and challenges obtaining homogeneous preparations of Nar1 in a fully Fe-loaded or a fully apo-state have led to persistent uncertainty regarding its molecular function of Nar1 in the CIA pathway.

One major barrier to resolving Nar1’s role is the lack of structural information and quantitative assays capable of defining how specific residues contribute to the Nar1-CTC interaction. Consequently, the molecular mechanism by which Nar1 engages the targeting complex and delivers its Fe–S cluster remains poorly understood. Here, we address this gap by combining affinity copurification experiments, quantitative fluorescence anisotropy binding assays and AlphaFold3 modeling to map the interaction interface between Nar1 and the targeting complex. Our results demonstrate that Nar1 engages the CTC through a bipartite interaction mechanism, with the primary site mediated predominantly by electrostatic interactions between a basic surface of Nar1 and an acidic surface lining the side of the third blade of Cia1’s β-propeller and a secondary site involving Nar1’s C-terminal tail engaging a shallow groove at the Cia1-Cia2 interface that comprises the TCR peptides binding site. Together these findings provide a mechanistic framework for understanding how Nar1 docks onto the targeting complex to enable Fe–S cluster delivery within the CIA pathway.

## MATERIALS AND METHODS

### Cloning

The plasmids for the heterologous expression of *C. thermophilum* (*Ct*) and *H. sapiens* (*Hs*) Nar1, with N-terminal His-SUMO tag, were created by insertion of the codon optimized gene into pTB146, analogous to heterologous overexpression plasmids for *Sc*Nar1.^37^ The plasmid for the heterologous expression of *H. sapiens* (*Hs*) double-tagged Cia1, with N-terminal His-TEV-Strep tag, were created by insertion of the codon optimized gene into pDUET1. Variants, including *Ct*Nar1^W609A^, *Hs*Nar1^Δ10^ (deletion of residues 462-471), and *Hs*Nar1^Δ15^ (deletion of 457-471) and *Hs*Nar1^ΔTCR^ (deletion of residues 472-476) were generated using Q5 Site-Directed Mutagenesis Kit (New England Biolabs). The SUMO peptide carrier (SPC) Nar1 was made by replacing nucleotides encoding the last 10 amino acids of *Ct*Nar1.^19,20^

The vector for co-expression of *Ct*Cia1-Cia2 and *Hs*Cia1 was previously reported.^19,20^ The *Ct*Cia1 charge-swapped variant (3E>K: E187K, E191K, E193K) and *Hs*Cia1 charge-swapped variant (D/E>K: D13K, E139K, E141K) were created by Q5 mutagenesis (New England Biolabs), using the appropriate co-expression plasmid as the template. Successful creation of all constructs was verified by DNA sequencing.

### Expression and Purification

Expression and purification of *Ct*Cia1-Cia2, *Hs*Cia1-Cia2a and SPC-Nar1 were carried out as described.^19,20^ Nar1 (*Ct* or *Hs* WT proteins or their variants) was expressed into BL21(DE3) grown at 37°C to an OD_600_ of 0.7−0.8. The temperature was then lowered to 30°C and IPTG (0.5 mM), ferric ammonium citrate (25 μM), and L-cysteine (2 mM) were added. The cells were collected 3.5-4 hours after IPTG addition.

For aerobic purification of *Ct*Nar1, cell paste (∼7g) was resuspended with 70 mL Buffer A [50 mM Na_2_HPO_4_ (pH 8.0), 100 mM NaCl, 5% glycerol, and 5 mM BME] supplemented with 0.5% octylthioglucoside (GoldBio), 5 mM Imidazole, 1 mM PMSF (GoldBio), protease inhibitor tablet (Thermo Scientific), 4 kU DNase nuclease (Thermo Scientific). rLysozyme (MilliPore Sigma) (45kU/g) was added and cells were disrupted at 4°C by sonication (20 s on/ 30 s off, 65% amplitude, 5 min). The clarified lysate was added to 1 mL of Ni-NTA resin which was washed with Buffer A supplemented with imidazole as follows, 40 column volumes (CV) with 5 mM imidazole, 20 CV with 10 mM imidazole, and 20 CV with 20 mM imidazole before elution of the column with Buffer A containing 300 mM imidazole. Protein containing fractions were combined and concentrated using Amicon ultra centrifugal filter units (10 kD MWCO, MilliporeSigma). Imidazole was removed by gel filtration using a PD10 column (Fisher Scientific) equilibrated in Buffer A. The protein was concentrated to 20-30 mg/mL and stored at -80 °C. As isolated, the protein contained 2.8±0.35 iron atoms and 2.6±0.54 acid labile sulfur per polypeptide (n=3).

For anaerobic purification of *Hs*Nar1, the cell pellet (∼6-8 g) was introduced into the glovebox (Coy Labs). Purification was carried out as described for the aerobic purification, except: cell lysis occurred via use of octylthioglucoside with no sonication and Buffer B [50 mM Tris (pH 8.5), 250 mM NaCl, 10% glycerol, and 5 mM BME] replaced Buffer A throughout the purification. A 2 mL Ni-NTA was used and *Hs*Nar1 was concentrated with an Amicon ultra centrifugal filter unit (30 kD MWCO, MilliporeSigma)). As isolated, the protein contained 3.7±1.23 iron atoms and 4.0±0.54 acid labile sulfur per polypeptide (n=3).

For aerobic purification of *Ct*Cia1-Cia2-Nar1 complex for SEC analysis, *Ct*Cia1-Cia2 and *Ct*Nar1 were separately expressed in BL21 (DE3) cells, grown at 37 °C to an OD_600_ of 0.7−0.8. After the temperature was shifted to 30 °C, expression was induced by adding IPTG (0.5 mM) along with ferric chloride (25 μM) and L-cysteine (2 mM). The cells were collected 3.5-4 hours after IPTG addition. For purification, cells overexpressing *Ct*Cia1-Cia2 (4 g) were mixed with those overexpressing *Ct*Nar1 (6 g). The resulting 10 g of the cell paste was resuspended in Buffer C [50 mM CHES (pH 9.0), 100 mM NaCl, 5% glycerol, 1 mM DTT; final volume 100 mL], supplemented with 1 mM PMSF, protease inhibitor tablet (Thermo Scientific), 4 kU DNase nuclease. Cells were treated with CellLytic Express (5.0 g). After 45 min incubation at 4°C with gentle stirring, the crude extract was centrifuged, and the soluble fraction was added to 2 mL Strep-Tactin XT Superflow resin (IBA). The column was washed with 15-20 CV of Buffer C and the protein was eluted with Buffer C supplemented with 30 mM D-Biotin (IBA) and 40 mM NaOH. Protein containing fractions were combined and concentrated using Amicon ultra centrifugal filter units (10 kD MWCO, Millipore Sigma), and the buffer was exchanged into 50 mM CHES pH 9, 100 mM NaCl, 2% glycerol, and 1 mM DTT by overnight dialysis. The protein concentration was determined using Assay Dye Reagent Concentrate (Bio-Rad) assuming a molecular weight of 154.5 kDa for the 1:1:1 complex, including tags. The protocol for isolation of *Ct*Cia1-Cia2 for SEC analysis was the same as the one described above for the *Ct*Cia1-CIa2-Nar1 complex but using the cell expressing *Ct*Cia1-Cia2 only.

For purification of *Hs*Cia1-Cia2a-Nar1, TEV protease treated *Hs*Cia1-Cia2a (TEV removes His-GST tag from *Hs*Cia2 and the His-tag from *His*Cia1, leaving a Streptactin tag on *Hs*Cia1 for affinity purification) was mixed with *Hs*Nar1 in a ratio of one μg *Hs*Cia1-Cia2a for every two µg *Hs*Nar1 in Buffer D [50 mM Hepes (pH 8.1), 100 mM NaCl, 5% glycerol]. After 1h at 4°C, the CCN complex was purified via streptactin chromatography.

### Size Exclusion Chromatography (SEC) analysis

All steps were carried out a 4°C. *Ct*Cia1-Cia2-Nar1 or *Ct*Cia1-Cia2 samples were diluted to 3.5 mg/mL in SEC-CHES buffer [50 mM CHES (pH 9.0), 100 mM NaCl, 2% glycerol, and 1 mM DTT]. The samples were centrifuged before 200-300 µL were loaded onto a Superdex 200 5/150 GL column (GE Healthcare) equilibrated with SEC-CHES buffer for *Ct* or SEC-HEPES buffer for *Hs* proteins. The column was eluted with at a 0.4 mL/min. The molecular weights were estimated by plotting the gel phase distribution coefficient (K_av_) against the logarithm of the molecular weight standards (Gel Filtration Standard, BioRad). Fractions (1 mL) were collected, concentrated to 200-240 µL using 10 kD Amicon ultra centrifugal filter units (MilliporeSigma) and analyzed by SDS-PAGE.

Samples of *Hs*Nar1-Cia1-Cia2a, *Hs*Cia1-Cia2a, and the individual subunits were diluted to 1-1.5 mg/mL and centrifugated to remove any aggregates. The samples (200 µL) were loaded onto a Superdex 200 5/150 GL column (GE Healthcare) equilibrated with 50 mM Hepes (pH 8.1), 100 mM NaCl, 5% glycerol, and 5 mM BME at 4°C and the column was eluted at a flow rate of 0.4 mL/min. Fractions (1 mL) were collected through the eluting peaks for subsequent analysis via SDS-PAGE.

### Affinity Copurification Analysis

Affinity copurification assays were carried out as previously described.^19,20^ Briefly, strep-tagged *Ct*Cia1-Cia2 complex (strep tag on Cia2) or strep-tagged *Ct*Cia*1* was mixed with an equimolar amount of *Ct*Nar1 in Buffer A (∼600-650 µL). After 1h incubation, the mixture was batch absorbed to Strep-Tactin XT Superflow (200 µL, IBA) resin. The resin was collected, washed, and eluted with Buffer A supplemented with 50 mM D-Biotin. The resulting elution fractions and the corresponding input samples were analyzed via SDS-PAGE. For the human orthologs, the experiments was performed similarly except *Hs*Cia1-Cia2a (Strep-tagged Cia1, 100-200 µg) and *Hs*Nar1 (200-300 µg) were mixed in a one-to-two mass ratio in 50 mM Hepes (pH 8.1), 100 mM NaCl, 5% glycerol.

### Fluorescence Anisotropy (FA) Binding Assay

Equilibrium binding assays with *Ct*Cia1-Cia2 complex and FITC-HLDW probe were carried out as described.^19^ Briefly, *Ct*Cia1-Cia2 (3 μM) or *Hs*Cia1-Cia2a (10 μM) was mixed with FITC-HLDW (0.05 μM) in the Buffer A (final volume 200 µL) with 0.1mg/mL BSA and unlabeled competitor (Nar1, 0-30 μM; SPC-Nar1, 0-150 μM) was added. The anisotropy values were recorded using a SpectraMax5 plate reader (MolecularDevices) In some experiments, the NaCl concentration was raised to 700 mM to determine how the ionic strength affects binding affinity.

### Bioinformatic analysis of Nar1’s TCR peptide

Eukaryotic sequences harboring the cytosolic Fe–S cluster assembly factor (IPR050340) were downloaded from InterPro. After removal of sequences shorter than 200 and longer than 1,000 amino acids and clustering of sequences into groups with ≥70% sequence homology and keeping one member of each group, the remaining sequences were aligned and manually processed to remove those missing any of the eight conserved cysteine residues or those not terminating in an aromatic residue. The last three residues of the remaining 631 Nar1 sequences were analyzed as described.^20^

## RESULTS

### Nar1 directly binds Cia1, and Cia2 further stabilizes formation of a defined three protein complex

Previous studies have reached conflicting conclusions regarding whether Nar1 is recruited to the CTC through its interaction with Cia1 alone, or whether Cia2 also plays a critical role in mediating the interaction.^19,25^ To resolve this discrepancy and direct comparison of interaction behavior across evolutionarily distant systems to arrive at a unified model, we examined interactions between Nar1, Cia1, and Cia2 using orthologs from fungi (*Chaetomium thermophilum, Ct*; *S. cerevisiae, Sc*) and humans (*Homo sapiens, Hs*). Throughout this manuscript, species-specific proteins are indicated with two letter prefixes (*Ct, Sc, Hs*), whereas generic protein names using the budding yeast nomenclature are used when referring to the protein family as a whole.

Whereas fungi encode a single Cia2 homolog, metazoans possess two, Cia2a and Cia2b.^11^ Here, we focused on human Cia2a based on its reported interaction with Nar1 in proteomic studies, its ability to form a ternary Cia1-Cia2a-Nar1 complex *in vitro* and its robust heterologous expression and purification, enabling direct biochemical comparisons across evolutionarily distant orthologs.^11,25,38^ The recombinant Nar1 proteins used in this study were purified with partially assembled Fe–S cofactors (approximately 3-4 Fe atoms per polypeptide; see *Materials* and *Methods* for species specific details), consistent with previous reports indicating incomplete cluster occupancy.^22,25,36^ In our experiments, the iron content corresponds to at most a single [4Fe-4S] cluster per Nar1 polypeptide. Attempts to generate fully apo preparations resulted in substantial destabilization, whereas efforts to increase cluster occupancy through chemical reconstitution or expression under conditions favoring Fe–S cluster assembly during heterologous overexpression did not yield substantially increased specific iron binding, as judged by the lack of increased spectroscopic signatures associated with effective incorporation of Fe–S clusters. We therefore performed the interaction studies using the as-isolated partially iron-loaded Nar1 preparations, which retained robust CTC binding activity.

Affinity copurification assays using the thermophilic fungal (*Ct*) and human (*Hs*) orthologs revealed that Nar1 associates with Cia1 in the absence of Cia2 (**Figure 2A, S1A-B**), corroborating our previous observations with the *S. cerevisiae* proteins.^19^ Notably, inclusion of Cia2 reproducibly increased Nar1 recovery across all orthologs (**Figure 2B** and **S1A-B**). In contrast, *Sc*Cia2 alone showed little to no interaction with *Sc*Nar1 and inclusion of *Sc*Met18 did not promote *Sc*Nar1 association with *Sc*Cia2 (**Figure S1E**). Thus, these data indicated that Cia1 is sufficient to bind Nar1, but Cia2 stabilizes the complex.

**Figure 2.**
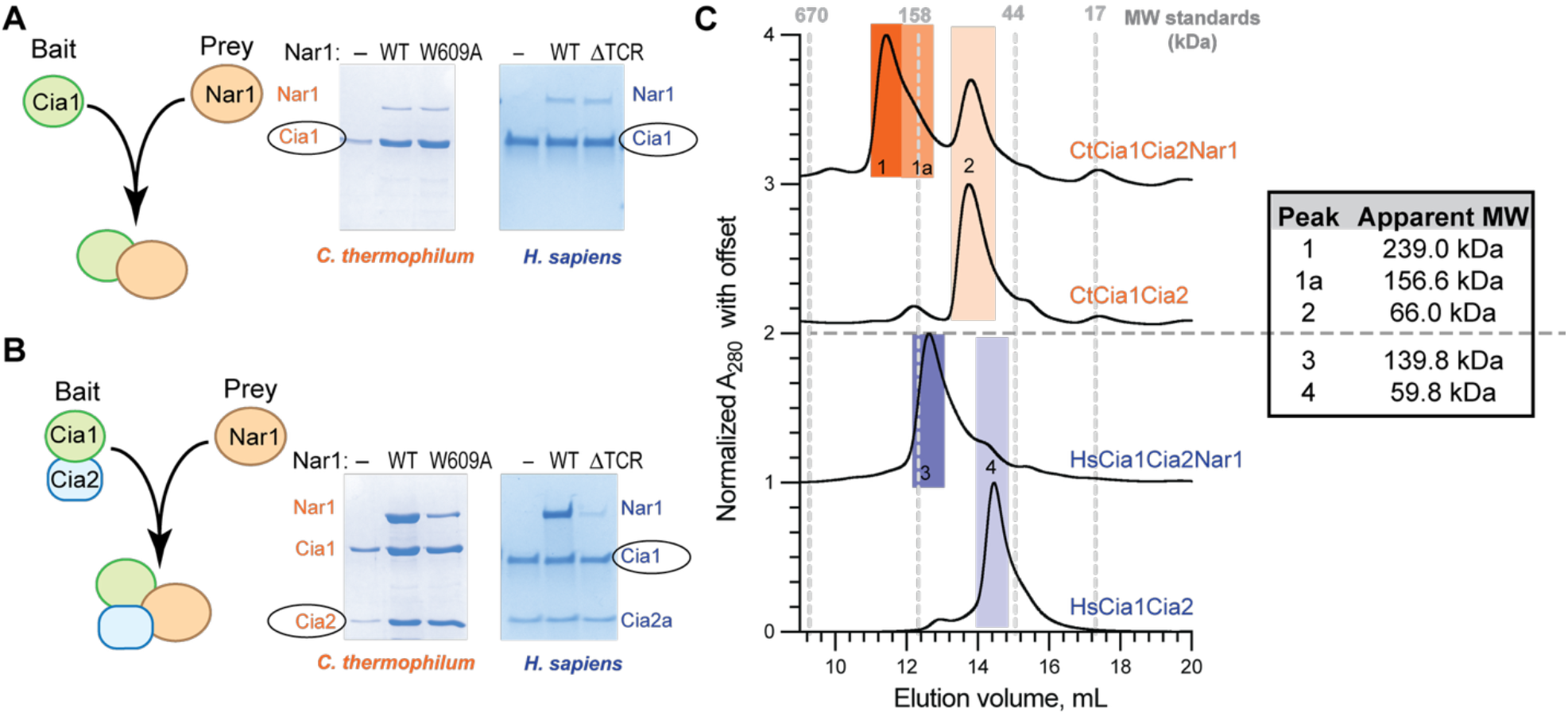
Interaction of Nar1 with Cia1-Cia2. **A-B**) SDS-PAGE analysis (Coomassie staining) of affinity copurification assays examining the interaction of Nar1 with Cia1 alone (**A**) or the Cia1-Cia2 complex (**B**). In each panel, the strep-tagged bait is circled. The Nar1 variant present in each lane is indicated above the gel as follows: no Nar1 (–), wild-type Nar1 (wt), *Ct*Nar1-W609A or *Hs*Nar1-ΔTCR (deletion of last five residues). Uncropped input and elution gels, including controls and molecular weight standards, are shown in **Figure S1**. (**C**) Size exclusion chromatography analysis of the indicated Cia1-Cia2 and Cia1-Cia2-Nar1 complexes. Elution volumes of molecular weight standards are indicated with dashed grey lines. Major peaks are indicated by numbered shaded boxes (*Ct*, orange, top); *Hs*, purple, bottom). The apparent molecular weight of major species is indicated to the right.

To probe the formation and stability of Cia1-Cia2-Nar1 (CCN) assemblies, we analyzed affinity copurified complexes by size exclusion chromatography (**Figure S2A-B**). *Ct*CCN eluted as a prominent high-molecular weight species (∼239 kDa) that was absent from the *Ct*Cia1-Cia2 control (**Figure 2C**). SDS-PAGE analysis confirmed the peak contained all three proteins (**Figure S2B**), indicating formation of a stable higher-order complex upon addition of *Ct*Nar1. Although the elution profile suggests the presence of assemblies larger than a simple heterotrimer, the shoulder (∼156 kDa) approximates the predicted molecular weight of a 1:1:1 Cia1:Cia2:Nar1 complex (155.1 kDa). Similarly, the human CCN complex eluted as a discrete high-molecular weight species (∼140 kDa) containing all three proteins (Figure 2C and **S2C-D**), consistent with formation of at least a heterotrimeric species (predicted molecular weight 122.1 kDa). These results indicate both the *C. thermophilum* and *H. sapiens* proteins can form stable CCN assemblies consistent with formation of heterotrimers as well as higher-order complexes (**Table S1**), aligning with previous reports of the ability of the CTC or its individual subunits to form higher-order assemblies.^17,18,37^ Together, these data demonstrate that Nar1 directly associates with Cia1 and Cia2 further stabilizes this three-protein complex, suggesting that Nar1 recruitment to the CTC involves contributions from both Cia1 and Cia2.

### Nar1’s divergent TCR motif recruits the CTC via a conserved pocket at the Cia1-Cia2 interface

Because our copurification studies indicated Nar1 recruits the CTC through a bipartite interaction involving both Cia1 and Cia2, we next sought to define the molecular features of this interface. Nar1 terminates in a conserved tryptophan that is essential for its role in the CIA pathway,^20^ reminiscent of the targeting complex recognition (TCR) motif found at the C-terminus of many CIA clients (**Figure 1A**). TCR peptides bind in a pocket formed at the Cia1-Cia2 interface and derive a substantial fraction of their binding energy from the terminal aromatic residue of this tripeptide motif. _19-21_

Despite this commonality, alignment of Nar1 orthologs strongly diverged from the [LIM]-[DES]-[FW] targeting peptide consensus. In contrast to CIA clients featuring an acidic residue in the penultimate position of the TCR motif, none of the 16 Nar1 sequences derived from diverse eukaryotic organisms matched this consensus (**Figure S3A**). To gain a comprehensive view of sequence conservation, we analyzed the C-terminal tripeptide of 631 Nar1 proteins. To our surprise, 45% of the sequences contained a basic lysine residue while only 4% had an acidic aspartate residue (**Figure S3B**). Likewise, the most common antepenultimate residue is a valine (23%), not a large aliphatic residue as seen in the canonical TCR motif. This divergence suggested either Nar1 does not engage the TCR peptide pocket or does so through a modified binding mode distinct from that used by canonical TCR peptides in CIA clients.

To test whether Nar1’s C-terminus contributes to CTC recruitment, we examined how perturbation of Nar1’s C-terminus impacts its association with Cia1-Cia2. Deletion of the final five residues of *Hs*Nar1 (ΔTCR) or substitution of the terminal Trp in *Ct*Nar1 (W609A) markedly reduced copurification with their respective Cia1-Cia2 complexes to levels comparable to Nar1 recovered with Cia1 alone (**Figure 2A-B** and **S1A-B**). In contrast, these variants had no detectable effect on the Nar1-Cia1 interaction in the absence of Cia2, indicating that Nar1’s C-terminal tail contributes to stabilization of the CCN complex through interactions involving Cia2.

To directly probe this interaction site, we substituted two conserved arginine residues within the TCR peptide binding pocket, one from Cia1 and one from Cia2, and assessed their impact on the formation of the CCN assembly.^19,21^ Substitution of the Cia2-derived residue (R177A-*Ct*Cia2; R138A-*Hs*Cia2a) impaired Nar1 recruitment, whereas mutation of the Cia1-derived arginine (R163A-*Ct*Cia1; R125A-*Hs*Cia1) had little to no impact (**Figure 3A-B** and **S1C-D**). This behavior contrasts sharply with CIA clients bearing canonical TCR motifs, which strongly depend on both arginine residues.^19^ These data indicate that Nar1’s C-terminus engages the TCR peptide binding pocket, likely through a non-canonical binding mode that depends strongly on Cia2, consistent with the divergence of Nar1’s tail from the TCR consensus.

**Figure 3.**
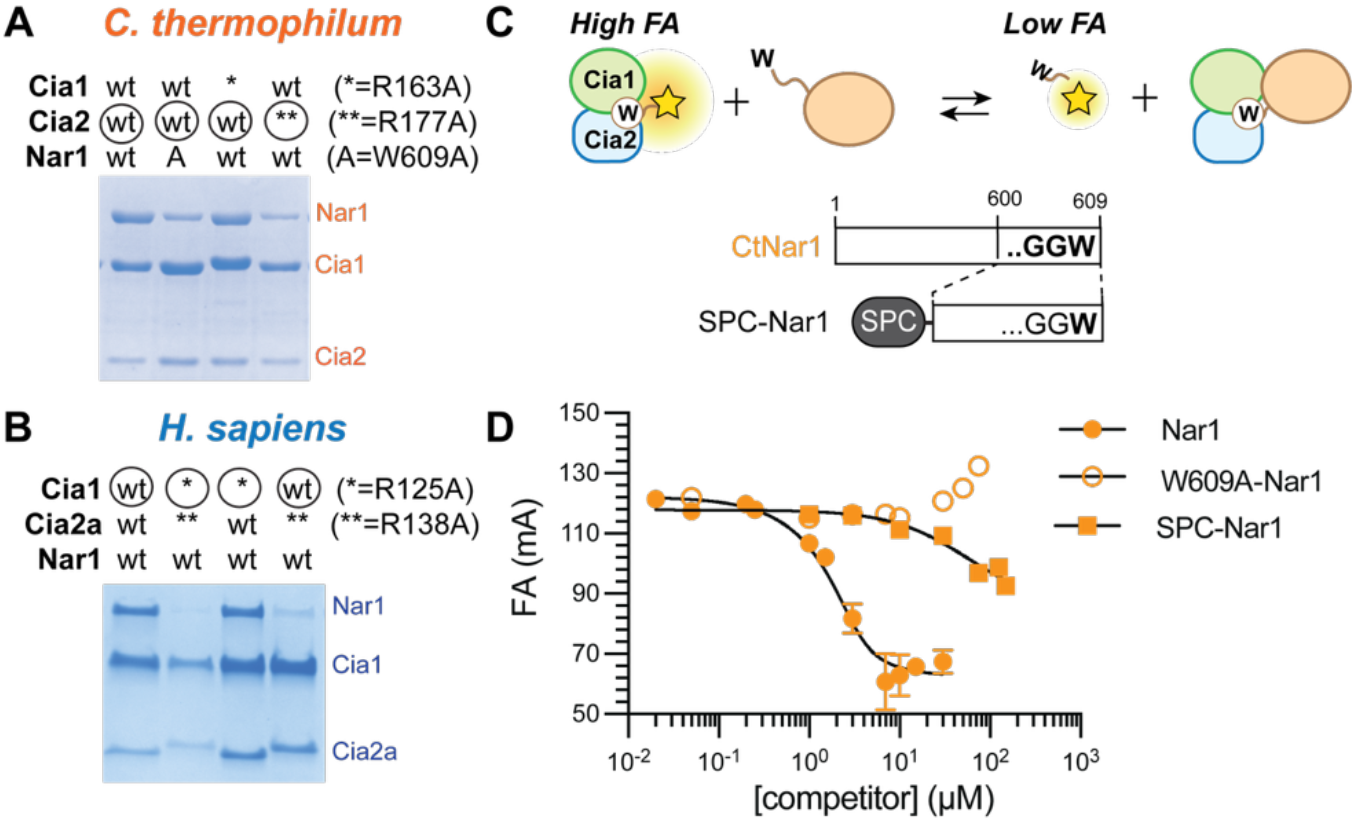
Interaction of Nar1 with the TCR peptide binding site of Cia1-Cia2. (**A-B**) SDS-PAGE analysis (Coomassie staining) of affinity copurification assays monitoring the interaction of Nar1 with variants of Cia1 and Cia2. Extended data, including inputs and controls, are provided in **Figure S1C-D**. (**C-D**) Scheme for the competitive fluorescence anisotropy (FA) assay (**C**) and the resulting data (**D**). Competitive FA assay in which the FITC-HLDW probe (0.05 μM) is displaced from *Ct*Cia1-Cia2 by unlabeled competitor, including wild-type *Ct*Nar1, *Ct*Nar1-W609A, or SPC-Nar1. The observed K_D_ values for *Ct*Nar1 and SPC-Nar1 were 0.4 µM and 190 µM, respectively. The W609A Nar1 variant did not displace the probe.

To quantify the energetic contribution of Nar1’s C-terminus, we performed fluorescence anisotropy (FA) competition assay using a FITC-labeled tetrapeptide probe (FITC-HLDW; **Figure 3C**).^19^ Full-length *Ct*Nar1 displaced the probe from *Ct*Cia1-Cia2 with an apparent dissociation constant (*K*_*D*_) of 0.4 µM, whereas the W609A variant failed to compete (**Figure 3D**). Similar qualitative results were observed with the human proteins (**Figure S4**), although protein aggregation precluded determination of a *K*_*D*_ value. To isolate the energetic contribution from the Nar1 tail alone, we fused the final ten residues of *Ct*Nar1 to SUMO, generating a SUMO peptide carrier for the *Ct*Nar1 C-terminus (SPC-Nar1).^19,20^ SPC-Nar1 displaced the peptide probe with an IC_50_ over two orders of magnitude higher than the full-length *Ct*Nar1 (**Figure 3D**). In total, the qualitative and quantitative binding data demonstrate that Nar1’s C-terminal tail is sufficient to engage the TCR peptide pocket but contributes only modestly to the overall binding affinity, consistent with the divergence of the Nar1 tail from the canonical TCR consensus. Thus, Nar1 must exploit an additional binding surface beyond the Cia2-dependent interaction mediated by its C-terminal tail.

### A conserved electrostatic interface anchors Nar1 to the Cia1 subunit of the CTC

Having established that Nar1’s C-terminal tail engages the TCR peptide binding pocket but contributes modestly to the overall binding affinity of the CCN complex, we next sought to identify the dominant interaction surface mediating the Cia2-independent interaction. Guided by AlphaFold models (*vide infra*), we first tested whether electrostatic or hydrophobic forces drive Nar1-Cia1 complex formation by examining Nar1 binding to the CTC under conditions of increased ionic strength. Increasing the NaCl concentration from 100 to 700 mM abolished detectable *Ct*Nar1-mediated displacement of the FITC-HLDW probe from *Ct*Cia1-Cia2 without affecting the binding of the FITC-labeled TCR peptide (**Figure 4A-B**). These data suggest that Nar1’s association with the CTC is strongly influenced by ionic interactions that are shielded as the buffer ionic strength is elevated.

**Figure 4.**
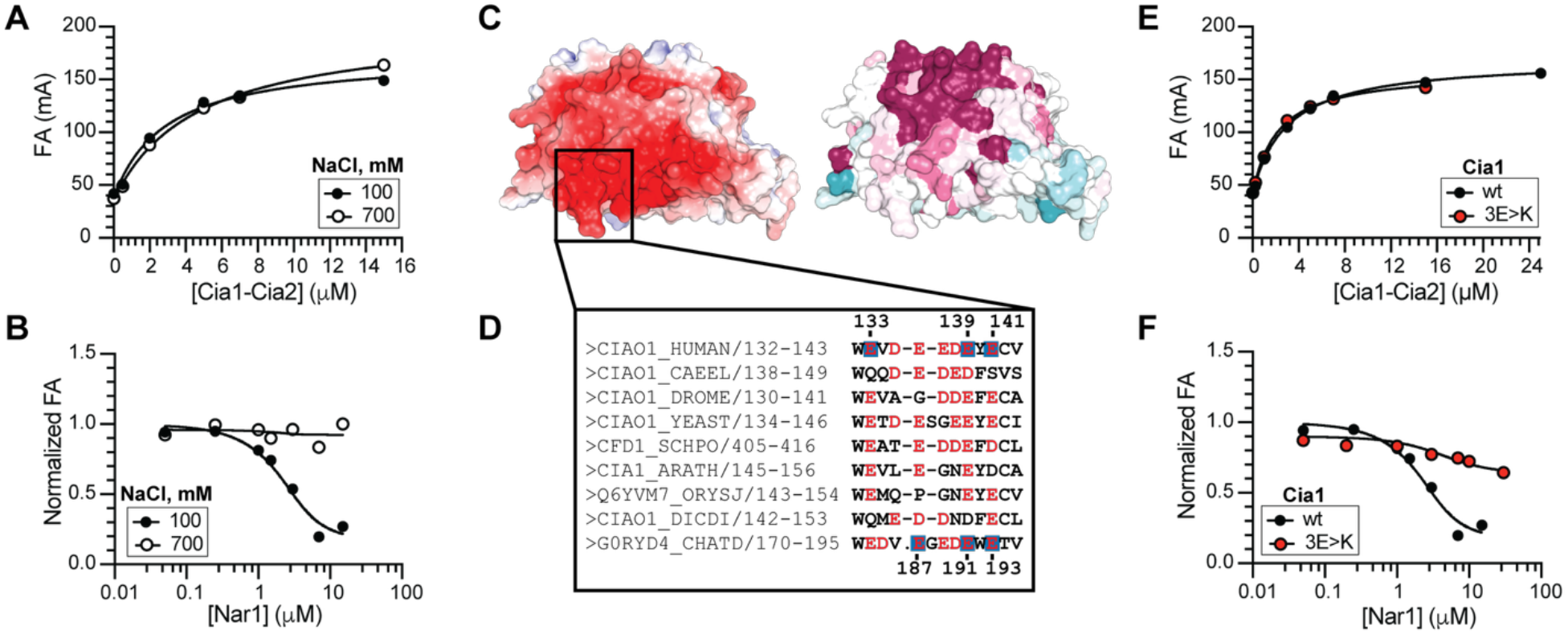
Nar1 docks on the side of Cia1 via a negatively charged interface. (**A**) FA assay showing FITC-HLDW probe (0.05 µM) binding to *Ct*Cia1-Cia2 (0-15 µM) under low (100 mM) and high (700 mM) concentrations of NaCl and establishing that the ionic strength does not impact TCR peptide binding affinity nor the integrity of the *Ct*Cia1-Cia2 complex. (**B**) Competitive FA assay in which the *Ct*Nar1 can displace the FITC-HLDW from *Ct*Cia1-Cia2 under low ionic strength (100 mM NaCl) conditions, but fails to do so when the ionic strength of the buffer is raised (700 mM NaCl). (**C**) Surface representations of *Hs*Cia1 (PDBID 3FMO)^39^ colored by electrostatic potential (left),^40,41^ or residue conservation (right; pink, conserved; cyan, variable)^42^ using the alignment in **Figure S5**. (**D**) Sequence alignment of Cia1 orthologs highlighting region of the Asp/Glu-to-Lys substitutions in the 3E>K variants (blue boxes, numbering in human (top) and *C. thermophilum* (bottom) are indicated). (**E**) FA assay (as in Panel A) showing that the *Ct*Cia1^3K>E^ variant (red) retains normal TCR peptide binding. (**F**) Competitive FA assay (as in Panel B) demonstrates that the charge-swapped variant (*Ct*Cia1^3K>E^, red) is strongly disrupted in *Ct*Nar1 binding to the *Ct*Cia1-Cia2 complex.

Guided by this result, we focused on a conserved acidic surface on blade 3 of the Cia1 β-propeller that has previously been implicated in CIA client binding (**Figure 4C** and **S5**).^17,43^ Because Nar1 and TCR-bearing clients compete for recruitment to the CTC,^19^ we hypothesized that this acidic patch may also serve the primary Cia1-centered docking site for Nar1. To test this, we replaced three conserved glutamate residues on *Ct*Cia1 (E187, E191, E193) with lysine, generating the charge-swapped variant *Ct*Cia1^3E>K^ (**Figure 4D**). This variant weakened *Ct*Nar1 binding to the *Ct*Cia1-Cia2 complex by ≥40-fold but did not affect Cia2 association nor TCR peptide binding affinity (**Figure 4E-F** and **S6A**; **Table S2**), indicating that these substitutions selectively disrupt Nar1 recruitment without affecting TCR peptide binding or the integrity of the Cia1-Cia2 complex as variants disrupting this complex are known to indirectly impact TCR peptide binding affinity. Analogous substitutions similarly weakened binding of *Hs*Cia1 to *Hs*Nar1 in affinity copurification assays (**Figure 4D** and **S6**), although the limited stability and solubility of the human proteins prevented determination of a dissociation constant. These quantitative and qualitative binding data identify the conserved, acidic patch on the side of blade 3 of Cia1 as a major contributor to Nar1 recruitment to the CTC, defining the second, Nar1-CTC interaction site that cooperates with Nar1’s C-terminal tail to stabilize CCN complex formation.

### AlphaFold Modeling of Nar1 and the CCN complex

Having defined two interfaces that work together to recruit the CTC to Nar1, we next used AlphaFold to generate structural models of Nar1 and the CCN complex because no experimental structures of Nar1 are available.^44^ These models allow visualization of the Nar1-CTC interfaces and evaluation of whether the predicted Fe–S binding sites of Nar1 and the targeting complex are spatially compatible with our biochemical data and with Nar1’s proposed role as an Fe–S cluster carrier for the CIA pathway.^1,12^

We first generated AlphaFold3 models of *Ct, Sc*, and *Hs* Nar1.^44^ All models revealed a well-folded ∼400-residue C-terminal domain and a lower-confidence N-terminal region (∼100 residues), consistent with the limited homology between Nar1 and [FeFe]-hydrogenase in this region (**Figure 5A, S7**, and **S8A**). Examination of the eight conserved Cys residues proposed to ligate Nar1’s Fe–S clusters (**Figure 1B-C**)^22^ revealed that the C-terminal cysteines form a defined pocket compatible with metallocluster coordination (**Figure 5A**). In contrast, the N-terminal cysteines were distributed within a low-confidence region that did not define a clear cofactor binding site.

**Figure 5.**
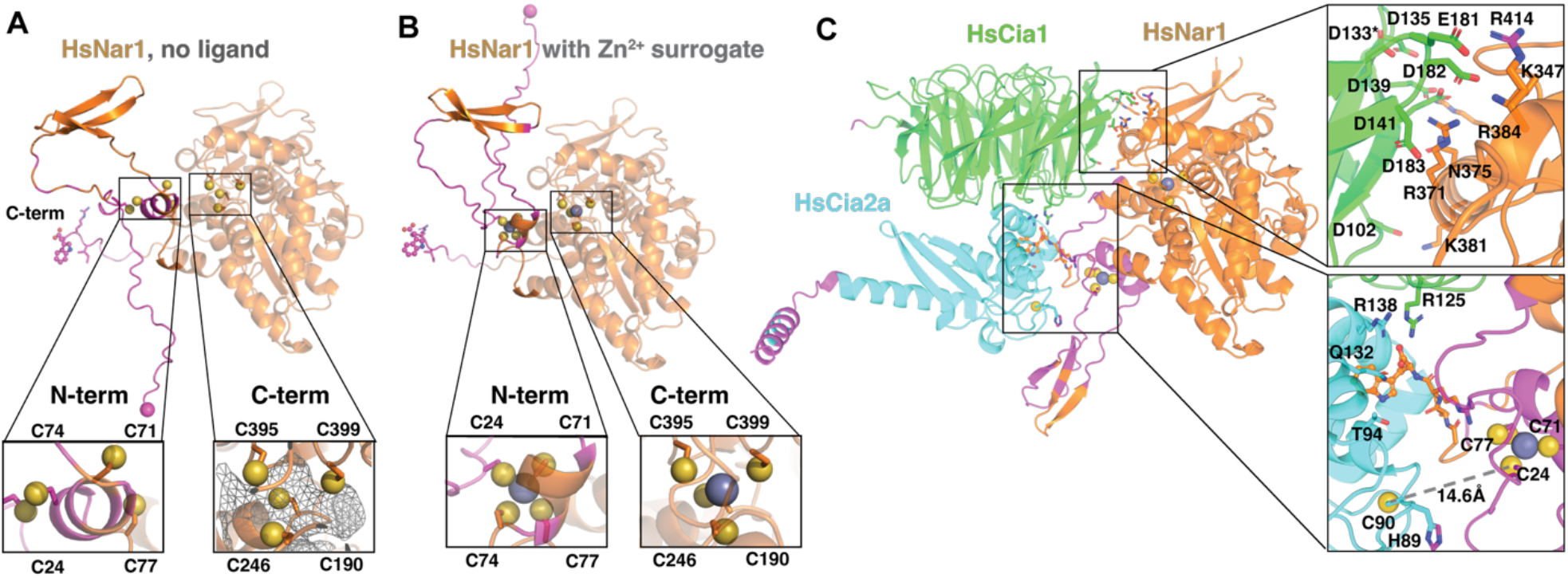
AlphaFold modeling of Nar1 and the CCN complex. A) Models of *Hs*Nar1 (orange), *Hs*Cia1 (green), *Hs*Cia2a (cyan), with regions of high uncertainty (pLDDT <50) in magenta. Cluster ligating Cys sidechains (yellow spheres), zinc ions (grey spheres), *Hs*Nar1 N-terminus (magenta sphere), and C-terminal TCR motif (sticks) are shown. Insets show detailed view of the N-terminal (N-term) and C-terminal (C-term) cluster binding cysteine residues. From left to right, models are *Hs*Nar1, *Hs*Nar1 with zinc, *Hs*Cia1-Nar1 with zinc, and *Hs*Cia1-Cia2a-Nar1 with zinc. B) and C) show the Cia1 (top) and Cia2a (bottom) surface facing Nar1. Two representations are shown, one colored by conservation (left) and the second by charge (right). D) shows the Hs Nar1 surface binding to Cia1 (above grey line) and Cia2 (below grey line) colored by charge (left) and conservation (right)

We reasoned that the absence of a bound cofactor might limit AlphaFold’s ability to organize this N-terminal region, as Fe–S clusters can serve as structural cores for small domains that lack extensive hydrophobic packing. Indeed, many Fe–S proteins, including cluster biogenesis factors,^45-47^ are well known to unfold in the absence of a bound Fe–S cluster. Because AlphaFold3 does not support modeling Fe–S clusters as ligands directly, we explored whether inclusion of other metal ions could stabilize this region and help locate the putative cluster binding sites. Although ferric iron can be specified as a ligand, its coordination chemistry differs substantially from that of Fe–S clusters. However, Zn^2+^ frequently occupies cysteine-rich coordination sites and Fe–S binding domains are sometimes misannotated as zinc fingers.^48^ These observations motivated us to include zinc ions as a Fe–S cluster surrogate. Indeed, inclusion of Zn^2+^ ions led to its coordination at both putative cluster binding sites and repositioned N-terminal cysteines into a geometry compatible with metal coordination (**Figure 5B**). However, neither inclusion of zinc nor Nar1’s binding partners (Cia1 and Cia2) resulted in folding of the N-terminal domain into a compact, globular domain (**Figure 5C** and **S8**). These observations suggest that the N-terminal region of Nar1 may be conformationally dynamic, consistent with a role of a cluster carrier requiring transient metallocluster binding during cytosolic iron sulfur cluster assembly.

Guided by our biochemical identification of a conserved acidic surface along blade 3 of the Cia1 β-propeller structure as a key Nar1 contacting region, we next examined the Nar1 models to identify a complementary interaction surface. All Nar1 orthologs displayed a prominent, basic region that is well conserved among Nar1 orthologs but absent in [FeFe] hydrogenases (**Figure S9**), consistent with a role in CTC recruitment. The predicted electrostatic complementarity between this basic patch on Nar1 and the acidic surface on Cia1 provides a structural rationale for the strong salt-dependence of the Nar1-Cia1 interaction observed experimentally (**Figure 4**).

Across all orthologs examined, AlphaFold3 predictions converged on similar CCN architecture, indicating that Nar1 docking geometry is conserved across evolutionary distant systems. These models predict that Nar1 engages the targeting complex through two interfaces: (1) the C-terminal tail of Nar1 engaging the TCR peptide binding pocket at the Cia1-Cia2 interface and (2) the basic surface of Nar1 contacting the acidic region of Cia1 blade 3 (**Figure 5C** and **S10**). Furthermore, the Cia1-Nar1 interface spans approximately 1100-1500Å^2^, consistent with our biochemical observation that Cia1 alone can recruit Nar1. By comparison, the TCR peptide binding site is substantially smaller, consistent with the mid-micromolar dissociation constant for the *Ct*Cia1-Cia2-SPC-Nar1 complex.

Notably, independently generated models using the Nar1 and Cia2 paralogs found in some animals, NarF and Cia2b respectively, also converged on similar CCN complex architectures (**Figure S8D-E**). Although, to our knowledge, there is no experimental evidence linking NarF to the CIA machinery, our observations that NarF maintains the eight proposed Fe– S binding cysteines, the basic patch, and the divergent TCR motif (**Figure S7**) indicates that it has retained the features required for CTC recruitment, consistent with the AlphFold model of the *Hs*Cia1-Cia2b-NarF complex. Additionally, the ability of Cia2b to facilitate Nar1 interaction is fully compatible with its proposed role as a general CIA targeting factor.^11^ To evaluate whether this CCN complex can be accommodated within the full CTC, we superimposed the *Hs*Cia1-Cia2a-Nar1 model on the published CTC crystal structure (PDB ID 6TC0).^17^ This comparison revealed that Nar1 can be accommodated within the monomeric assembly but would sterically clash with the dimeric arrangement observed in the crystal lattice (**Figure S8F**). This observation is consistent with previous structural studies indicating that the monomeric CTC, containing one copy each of Cia1, Cia2 and Met18, represents the physiologically relevant complex for client recognition.^17^

We noted that in all of the CCN models the flexible N-terminal region of Nar1 is positioned near the proposed Fe–S acceptor site, defined by His89 and Cys90 (*Hs*Cia2a numbering; **Figure 5C**).^49,50^ Although these models and our biochemical studies herein do not directly address the mechanism of metallocluster transfer, the spatial proximity between the predicted Nar1 N-terminal cluster binding cysteine residues and the proposed cluster accepting residue of Cia2 is compatible with a model in which bipartite Nar1 docking positions the predicted cluster donor and acceptor sites in an orientation that would permit cluster transfer.

To probe the importance of this cooperative bipartite docking, we altered the relative positioning of *Hs*Nar1’s two CTC interaction sites, shortening the region connecting TCR motif to its globular C-terminal domain. These truncations markedly reduced recovery of *Hs*Nar1 with *Hs*Cia1-Cia2 in pulldown assays (**Figure S11**), indicating that the precise three-dimensional arrangement of these two CTC binding sites facilitates productive Nar1 binding to the targeting complex. Together, the models provide a structural framework that integrates our biochemical observations and supports a model in which Nar1 engages the CTC via two distinct binding interfaces that collaborate to stably bind Nar1 to the CTC. This bipartite interaction anchors Nar1 to the targeting complex while orienting its conformationally flexible N-terminal region toward the cluster binding surface of Cia2, providing a plausible structural arrangement for productive engagement of the CIA targeting complex during cytosolic Fe–S protein maturation.

## DISCUSSION

In this study, we define the molecular basis by which Nar1 (also known as CIAO3/IOP1/NARFL) engages the CIA targeting complex (CTC) through a bipartite binding mechanism.^1,19,25,30^ Using cross-species comparisons together with qualitative and quantitative binding analyses, we demonstrate that Nar1 is recruited through two distinct interfaces: a primary, predominantly electrostatic interaction that anchors Nar1 to the conserved acidic surface lining Cia1’s third β-propeller domain,^16,17^ and a secondary interaction in which Nar1’s C-terminal tail, bearing a divergent targeting complex recognition (TCR) motif, engages a peptide binding pocket at the Cia1-Cia2 interface.^19^ Together with AlphaFold-based modeling, these data provide a structural framework that reconciles prior conflicting models for Nar1 recruitment and explains how formation of the Cia1-Cia2-Nar1 (CCN) complex positions the N-terminal Fe–S binding domain of Nar1 adjacent to the CTC cluster binding site.

A primary outcome of this work is resolution of longstanding discrepancies regarding the minimal requirements for Nar1’s recruitment to the targeting complex. Earlier studies suggested Nar1 interacts primarily with Cia1,^30^ whereas later work concluded that the assembled Cia1-Cia2 complex is required.^25^ Our results reconcile these contradictory conclusions by showing that Cia1 provides the principal recruitment interface, while Cia2 enhances affinity through creation of the secondary Nar1 binding site situated at the Cia1-Cia2 interface. This cooperative mechanism ensures that stable Nar1 association only after formation of a functional CTC core, comprising Cia1 and Cia2, preventing premature interaction with isolated Cia1 which lacks Fe–S cluster binding residues. Notably, modeled CCN assemblies presented herein position the Nar1 N-terminal Fe–S cluster binding domain adjacent to the conserved, reactive Cia2 cysteine residue (Cys90 of *Hs*Cia2a).^15,49^ This quaternary structure is consistent with previous studies indicating that the N-terminal region of Nar1 is consistent with proposals that the N-terminal region participates in cluster trafficking.^22,25^ We speculate this multi-partite interaction serves to properly orient the putative Fe–S donor and acceptor sites within the CCN complex in a manner that is compatible with cluster transfer.^17,20^

The structural models further suggest that Nar1 represents an evolutionary repurposing of a hydrogenase-like scaffold for a metallochaperone function in the CIA system. Although Nar1 retains cysteine motifs characteristic of [FeFe]-hydrogenases, it lacks the catalytic H-cluster and instead has acquired new features, such as the basic patch implicated in CTC recruitment. The absence of this electrostatic surface in [FeFe]-hydrogenases suggests Nar1 has acquired this new protein-protein interaction module to facilitate its role as a trafficking factor with the CIA pathway.

Our findings further demonstrate that Nar1 engages a conserved, client-recruitment interface of the CTC. Previous structural studies implicated a negatively charged surface along the side of Cia1’s third β-propeller domain in facilitating client recognition.^16,17^ Our biochemical analysis pinpoints this same region, the dominant, Cia2-independent interaction site for Nar1 recruitment. Disruption of this acidic surface selectively weakens CCN formation without impacting Cia2 binding or TCR peptide recognition, indicating that it functions as an organizational hub coordinating interactions with the Nar1 cluster donor and CIA client protein acceptors. The use of shared docking interfaces to coordinate interactions with multiple binding partners represents a recurring organizational principle in Fe–S cluster biogenesis pathways. For example, the bacterial cysteine desulfurase IscS engages its various sulfur acceptors IscU and TusS via overlapping binding sites.^51,52^ Similarly, frataxin and ferredoxin compete for binding to IscU, tuning the rate of Fe–S cluster assembly by the ISC scaffolding complex.^53,54^ The parallels suggest that competitive engagement of a common recruitment interface might similarly function in CIA to coordinate exchange of an Fe–S cluster between the CTC and the Nar1 donor and a downstream CIA client acceptor. Together with previous studies identifying this same surface in client recognition, these findings suggest that this region of the CTC functions as a central recruitment hub that coordinates sequential interactions with cluster donors and client proteins.

Bioinformatic analyses of Nar1 and [FeFe]-hydrogenases pinpointed Nar1’s acquisition of a TCR peptide tail as a second functionally important divergence between Nar1 and hydrogenase proteins. We reveal that Nar1 engages the same TCR peptide binding client pocket exploited by CIA clients with canonical TCR motifs, but through a noncanonical interaction. Bioinformatic analysis reveals that the C-termini of Nar1 diverge from those of clients bearing a canonical TCR motif primarily at the penultimate position, where an acidic residue is typical in CIA clients is frequently replaced by a basic residue and the antepenultimate bulky aliphatic residue is replaced with the small valine. Since the CTC binds Nar1 and clients through a shared interface, we speculate the divergence of their TCR motifs may allow the targeting complex to distinguish between Fe–S cluster donors and client acceptors. Consistent with this divergence, the recognition of Nar1’s C-terminus depends predominantly on the Cia2 arginine hotspot residue rather than contributions from both Cia1 and Cia2 as required for canonical TCR peptide binding.^19^ The proximity of the TCR peptide binding pocket to the proposed Fe–S binding site on Cia2 further suggests that TCR peptide engagement may influence cluster transfer.

A major barrier to investigations of cluster transfer between Nar1 and the CTC is the difficultly isolating Nar1 in a defined state as previous cell based studies have found the interactions investigated herein are regulated by iron.^24^ As isolated, the Nar1 preparations used herein contain 3-4 irons per polypeptide, in agreement with work by others.^22,25,36^ In an attempt to create fully apo-forms of Nar1, we attempted to mutate the Fe–S cluster binding cysteines but found that deletion of just one ligand did not prevent iron association and, consistent with the reports of others, additional substitutions disrupted Nar1 folding and thus precluded use in biochemical investigations.^22,25^ Similarly, attempts to reconstitute Nar1 via chemical reconstitution resulted in incomplete cluster incorporation. We note that it was recently reported that *Sc*Nar1 could be reconstituted with two [4Fe4S] clusters and one [2Fe2S] cluster, but the ability of Nar1 to donate one or more of these clusters to the CTC was not investigated in that or any system.

The bipartite recruitment mechanism defined herein provides a structural framework for understanding how Nar1 engages the CIA targeting complex and lays the foundation for future mechanistic studies investigating how the CTC regulates interaction with the Nar1 donor and the CIA client acceptors during cytosolic iron–sulfur protein maturation. A key remaining challenge is to obtain clear in vitro evidence for Nar1’s proposed Fe–S donation activity and, more broadly, how iron modulates the complex network of interactions in the CIA pathway.^24^ Given the emerging links between defects in cytosolic Fe–S biogenesis and human disease,^6,7,9,14^ defining how cluster exchange occurs within the CTC-Nar1 complex represents an important next step to understanding how its disruption step contributes to cellular dysfunction.

## Acknowledgement

This work was supported by the NSF-GRFP (M.M.), the NIH R01GM 121673 (D.L.P), and by the Boston University Undergraduate Research Opportunities Program (B.W.).

## Notes

### Competing Interest Statement

The authors have declared no competing interest.

