## Supporting Information for "Nar1 binds the cytosolic iron sulfur cluster assembly targeting complex via a bipartite interaction interface"

Includes:

Supporting Tables S1-S2

Supporting Figures S1-S11

**Table S1. Size exclusion chromatography data for CCN complexes**

| Organism | Sample | Peak | $V_e$ (mL) | Apparent MW (kDa) | Proposed Quaternary Structure, theoretical MW (kDa)[1] |
|---|---|---|---|---|---|
| *Hs* | Cia1 | - | 15.08 | 44.8 | Cia1, 41.8 |
| | ΔN27-Cia2a | - | 16.46 | 23.6 | Cia2a, 15.5 |
| | Nar1 | - | 13.95 | 75.6 | Nar1; 67.2 |
| | Cia1-Cia2a | - | 14.46 | 59.8 | Cia1-Cia2a; 54.9 |
| | Cia1-Cia2a-Nar1 | - | 12.63 | 139.8 | Cia1-Cia2a-Nar1, 122.1;<br>Cia1-Cia2a$_2$-Nar1, 137.6 |
| *Ct* | Cia1-Cia2 | 1 | 12.18 | 159.0 | (Cia1-Cia2)$_2$, 151.2 |
| | | 2 | 13.75 | 68.1 | Cia1-Cia2, 75.6 |
| | Cia1-Cia2-Nar1 | 1 | 11.43 | 239 | (Cia1-Cia2)$_2$-Nar1, 230.7<br>Cia1-Cia2-Nar1$_2$, 234.6 |
| | | 1a | 12.21 | 156.6 | Cia1-Cia2-Nar1, 155.1 |
| | | 2 | 13.81 | 66.0 | Cia1-Cia2, 75.6 |

[1]The following are the predicted molecular weights of monomeric proteins, accounting for any purification tags: *Hs*Cia1 (with 6xHisTEVStrep), 41.8 kDa; ΔN27-*Hs*Cia2a (untagged), 15.5 kDa; *Hs*Nar1 (with 6xHisSUMO), 67.2 kDa; *Ct*Cia1 (with 6xHis-TEV tag), 51.4 kDa: *Ct*Cia2 (with Strep-TEV tag), 24.2 kDa; *Ct*Nar1 (with 6xHis-SUMO tag), 79.5 kDa.

**Table S2.** Apparent dissociation constants related to **Figure 3 and 4.**

| Assay Format | Receptor | Ligand | [NaCl], mM | $K_D$, μM |
|---|---|---|---|---|
| **DIRECT BINDING ASSAY** 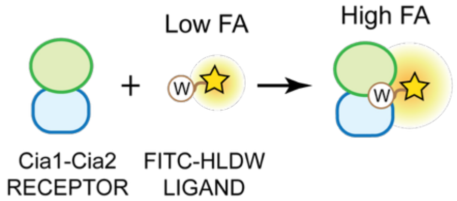  | WT *Ct*Cia1-Cia2 | FITC-HLDW | 100 | 2.5±0.5 (n=4) [1] |
| | WT *Ct*Cia1-Cia2 | FITC-HLDW | 700 | 3.2 (n=1) |
| | *Ct*Cia1$^{3E>K}$-Cia2 | FITC-HLDW | 100 | 3.7 (n=1) |
| **COMPETITIVE BINDING ASSAY** 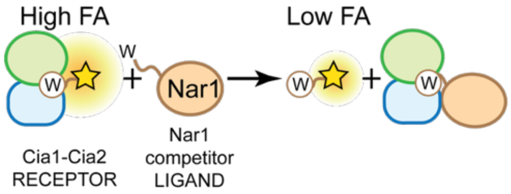  | WT *Ct*Cia1-Cia2 | WT *Ct*Nar1 | 100 | 0.4±0.2 (n=3) [2] |
| | WT *Ct*Cia1-Cia2 | WT *Ct*Nar1 | 700 | >200 (n=1) |
| | WT *Ct*Cia1-Cia2 | *Ct*Nar1-W609A | 100 | >200 (n=1) |
| | WT *Ct*Cia1-Cia2 | SPC-Nar1 | 100 | ≥180 (n=1) |
| | *Ct*Cia1$^{3E>K}$-Cia2 | WT *Ct*Nar1 | 100 | 43 (n=1) |

[1] The apparent dissociation constant of Cia1-Cia2 variants toward FITC-tetrapeptide was measured via the direct binding assay and analyzed as described,[19] where maximum anisotropy value was constrained to 180, the value observed with WT in the assay buffer with 100 mM NaCl.

[2] The apparent dissociation constant of Cia1-Cia2 variants to Nar1 variants was measured via the competition FA assay and data was fitted to quadratic equitation with [P] = 3 μM.

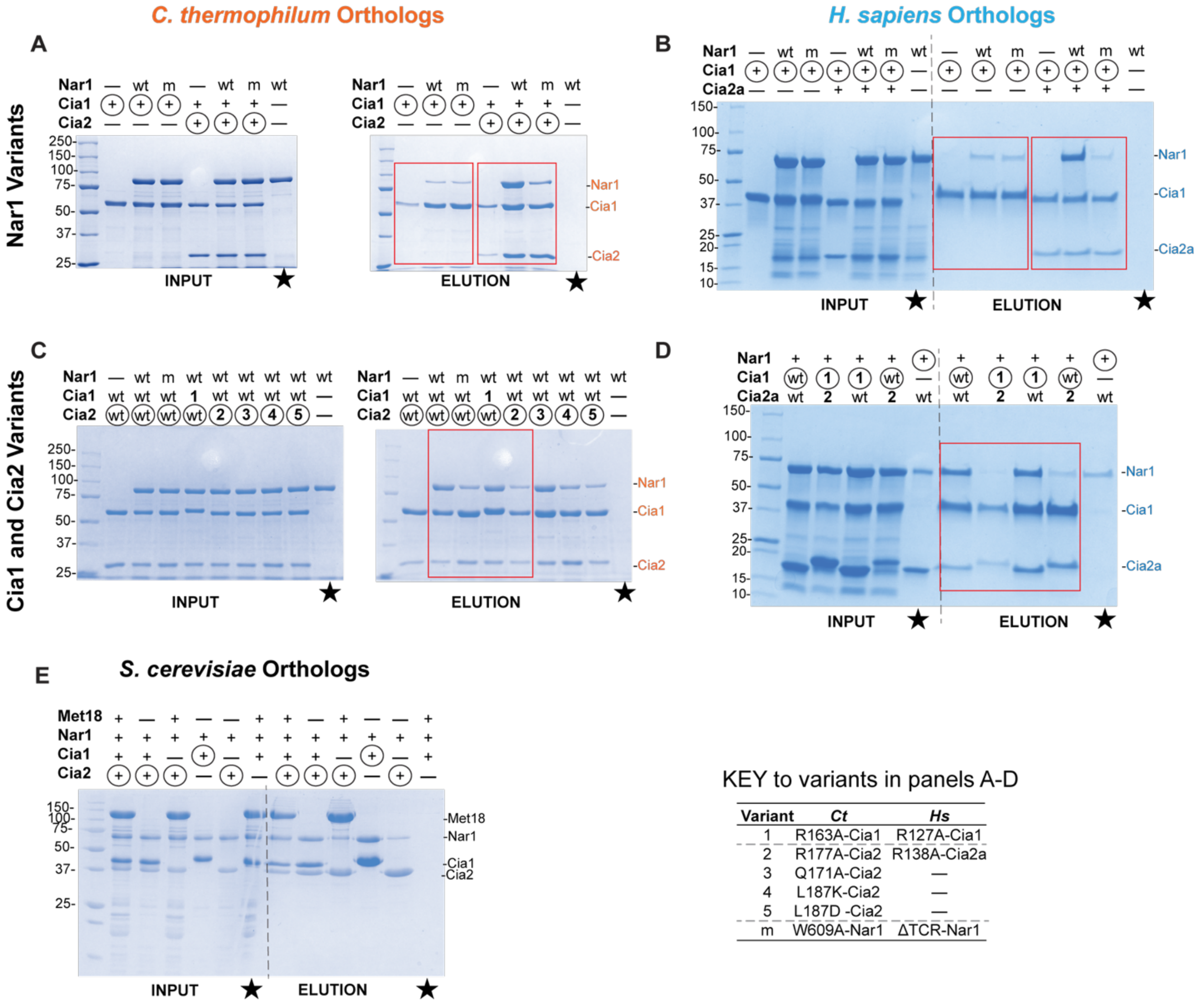


KEY to variants in panels A-D

| Variant | *Ct* | *Hs* |
|---|---|---|
| 1 | R163A-Cia1 | R127A-Cia1 |
| 2 | R177A-Cia2 | R138A-Cia2a |
| 3 | Q171A-Cia2 | — |
| 4 | L187K-Cia2 | — |
| 5 | L187D -Cia2 | — |
| m | W609A-Nar1 | ΔTCR-Nar1 |

**Figure S1. Nar1 interaction with the CTC assessed via affinity copurification assays, related to Figure 2 and 3.** Uncropped SDS-PAGE gels (Coomassie staining) showing input samples (before streptactin column) and streptactin affinity purification elution samples used to assess Nar1 association with Cia1 and Cia2 complexes from *C. thermophilium* (*Ct*, **A**, **C**), *H. sapiens* (*Hs*, **B**, **D**), and *S. cerevisiae* (*Sc*, **E**). Wild-type Nar1 or the C-terminal TCR mutants (m, *Ct*Nar1-W609A or *Hs*Nar1-ΔTCR) were mixed with the indicated combinations of Cia1 and Cia2 prior to purification using streptactin-tagged bait protein (circled). Panels **A** and **B** evaluate the contribution of the Nar1 C-terminal tail to binding of *Ct*Cia1-Cia2 and *Hs*Cia1-Cia2 complexes, respectively. Panels **C** and **D** assess the importance of conserved TCR peptide-binding residues on Cia1 and Cia2, using the variants indicated in the key. Regions corresponding to gels shown in the main text figures are boxed in red. Control purifications lacking a strep-tagged bait protein are indicated with the star.

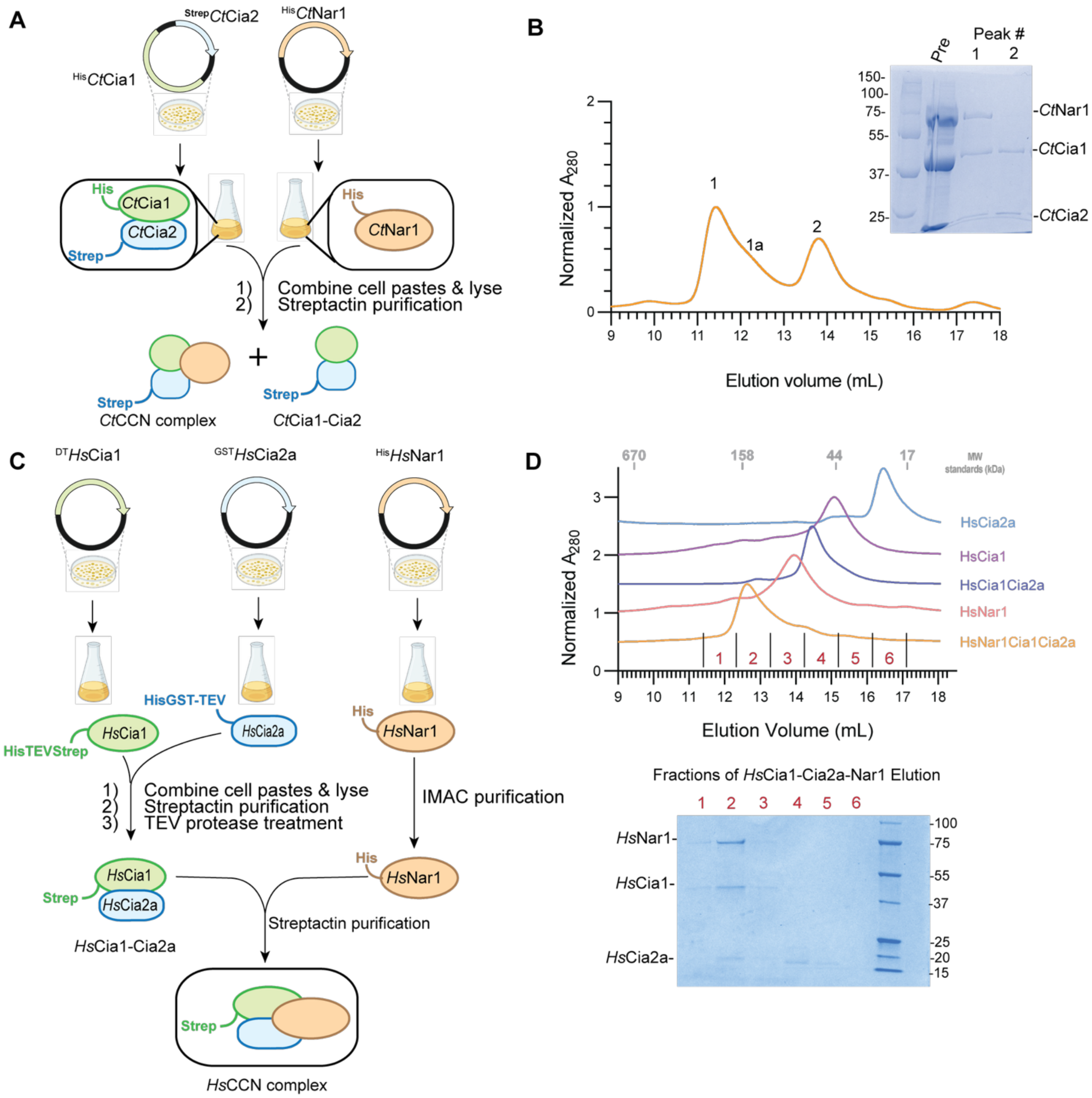


**Figure S2. Copurification and Size-exclusion chromatography analysis of Cia1-Cia2-Nar1 (CCN) complexes, related to Figure 2. (A)** A schematic illustrating purification of the *Ct*CCN complex. Strep-tagged *Ct*Cia1-Cia2 (coexpressed from a single plasmid) and *Ct*Nar1 were expressed and the resulting cell pastes were combined, allowing for *Ct*CCN assembly prior to streptactin purification. (**B**) SEC profile of the purified *Ct*CCN complex with corresponding SDS-PAGE analysis of elution peaks. The primary peak (**1**; apparent molecular weight 239.0 kDa) and its shoulder (**1a**, 156.6 kDa) contain *Ct*Cia1, *Ct*Cia2 and *Ct*Nar1, consistent with formation of a heterotrimer (Peak 1a) and higher order assemblies. Peak 2 (66 kDa) corresponds to *Ct*Cia1-Cia2 complex. Calculated molecular weights and proposed quaternary structures are summarized in **Table S1**. (**C**) Schematic of *Hs*CCN complex purification. *Hs*Cia1 and *Hs*Cia2a, with the indicated tags, were copurified and treated with TEV protease. The resulting complex was combined with *Hs*Nar1 and the resulting *Hs*CCN complex was purified via streptactin purification. (**D**) SEC analysis of individual human CIA components and assemblies as indicated. Elution of the *Hs*CCN complex (orange trace) results in co-elution of all three proteins within the primary peak (Fraction 2, red), confirming formation of the CCN complex. Calculated molecular weight and proposed assemblies are summarized in **Table S1**.

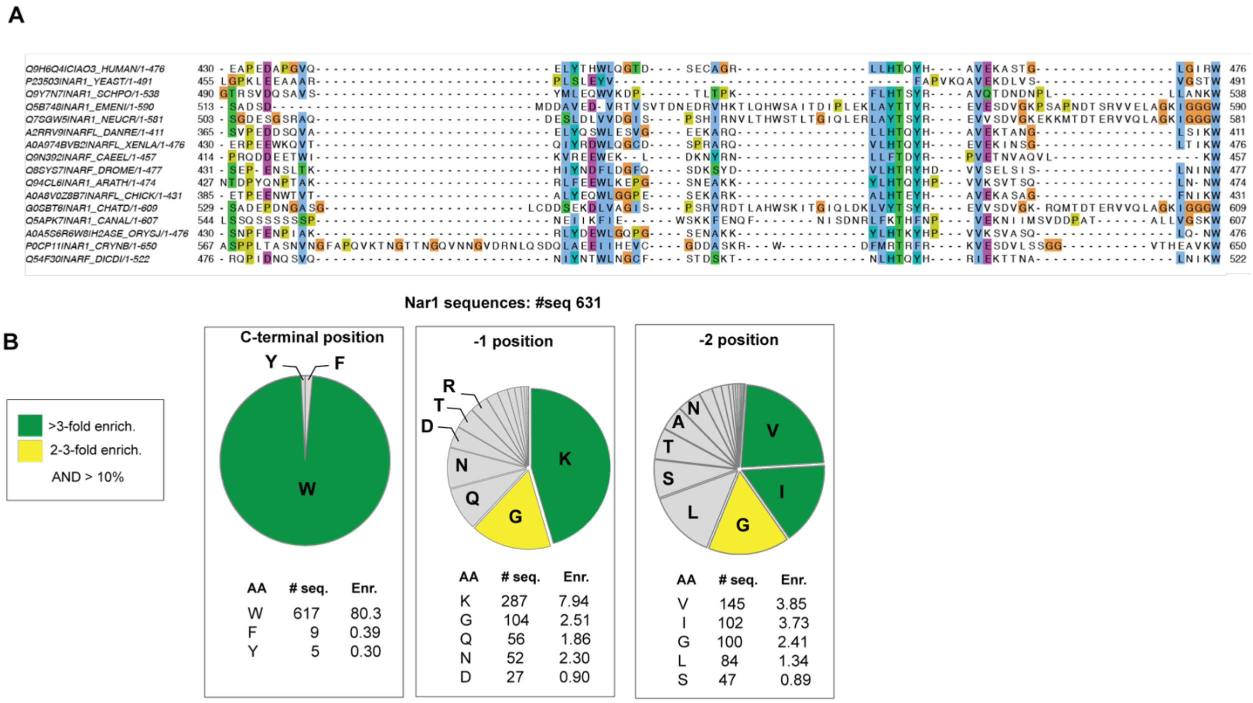


**Figure S3. Bioinformatic analysis of Nar1 C-terminal TCR motif.** (**A**) Multiple sequence alignment of representative Nar1 homologs highlighting conservation of the C-terminal TCR motif. The organisms and Uniprot accession numbers are as follows: *H. sapiens*, CIAO3_HUMAN; *S. cerevisiae*, NAR1_YEAST; *S. pombe*, NAR1_SCHPO; *E. nidulans*, NAR1_EMENI; *N. crassa*, NAR1_NEUCR; *D. rerio*, NARFL_DANRE; *X. laevis*, A0A974BVB2_XENLA; *C. elegans*, NARF_CAEEL; *D. melanogaster*, NARF_DROME; *A. thaliana*, NAR1_ARATH; *G. gallus*, A0A8V0Z8B7_CHICK; *C. thermophilum*, G0SBT6_CHATD, *C. albicans,* NAR1_CANAL; *O. sativa*, A0A5S6R6W8_ORYSJ; *D. discoideum*, NARF_DICDI; *C. neoformans*, NAR1_CRYNB. (**B**) Amino acid consensus sequence analysis for Nar1 C-terminal tripeptides (n=631 sequences, see Materials and Methods for description of the sequence set used in the analysis). The number of sequences containing each residue at the indicated position is shown together with enrichment values calculated as the frequency normalized by the frequency of that amino acid in the human proteome.[19] Residues present in ≥10% of sequences and enriched between 2-3-fold or >3-fold are indicated in yellow and green, respectively.

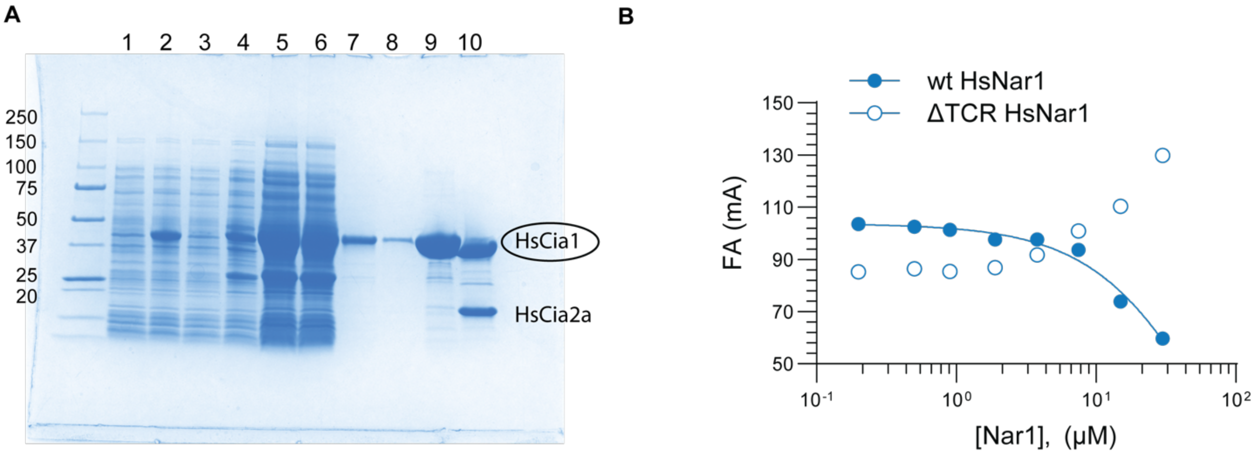


**Figure S4. The divergent TCR motif of *Hs*Nar1 is essential for its interaction with *Hs*Cia1-Cia2a. (A)** SDS-PAGE analysis (Coomassie staining) showing copurification of *Hs*Cia1-Cia2a. Cell lysates expressing doubly tagged (DT) *Hs*Cia1 (His-TEV-Strep; 41.8 kDa) and *Hs*Cia2a (His-GST-TEV; 43.7 kDa) were combined prior to lysis and purified by streptactin copurification as described in Figure **S2C**. Lanes are as follows: Lane 1 and 2, *Hs*Cia1 before and after IPTG addition, respectively; Lane 3 and 4, *Hs*Cia2a before and after IPTG addition, respectively; Lane 5, soluble lysate; Lane 6, streptactin column flow through; Lane 7-8, streptactin wash fractions; Lane 9, streptactin elution fraction; Lane 10, TEV protease treated and purified *Hs*Cia1-Cia2a complex. **(B)** Fluorescence anisotropy (FA) competition monitoring displacement of FITC-LDW peptide (0.05 μM) from *Hs*Cia1-Cia2a (10 μM) by wild-type *Hs*Nar1 or the ΔTCR variant. Wild-type Nar1 competes efficiently for binding, whereas the ΔTCR variant does not. Increased anisotropy observed at high concentrations of *Hs*Nar1-ΔTCR is attributed to aggregation that interferes with the spectroscopy.

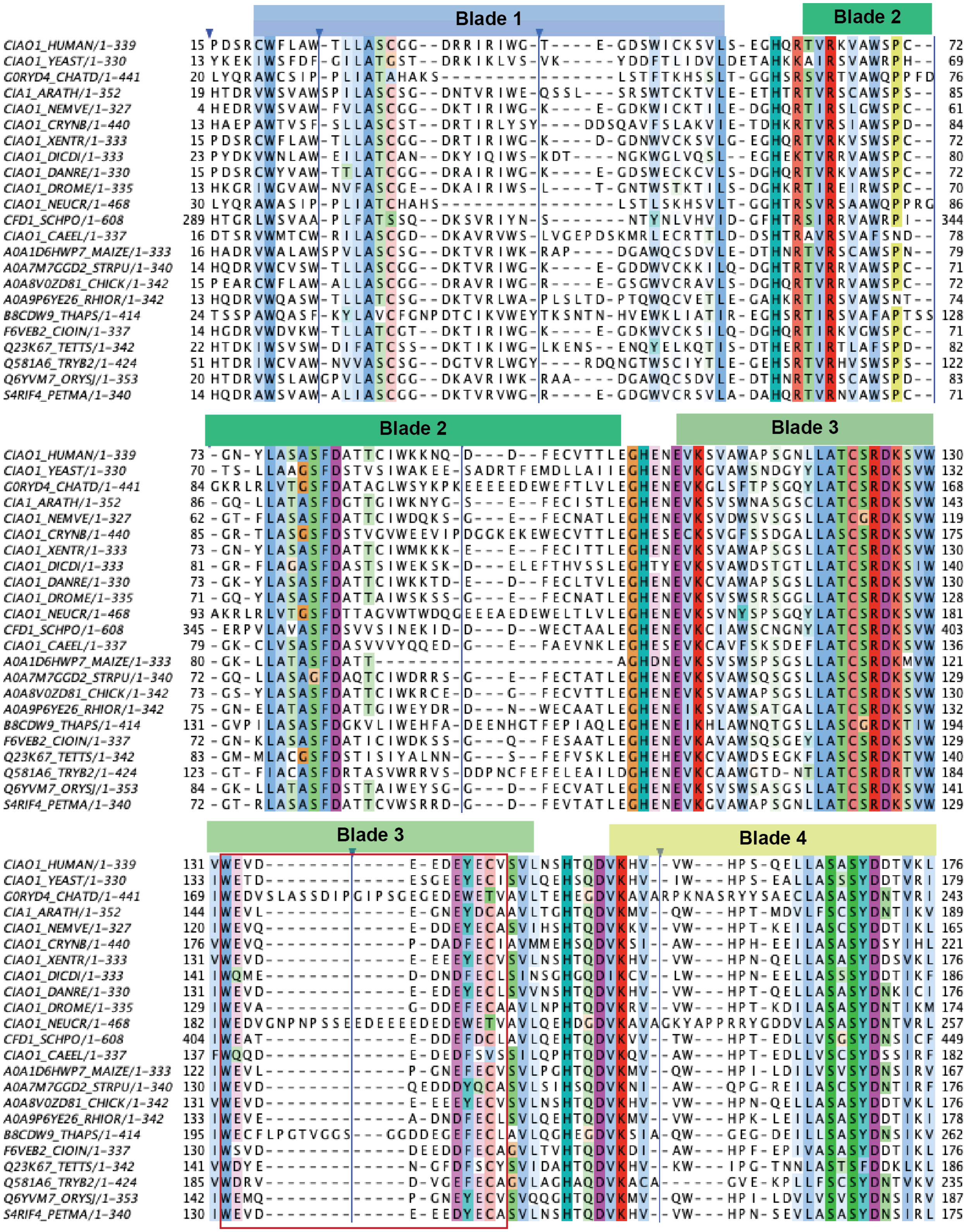

Blade 1
Blade 2
Blade 3
Blade 4
CIAO1_HUMAN/1-339
CIAO1_YEAST/1-330
G0RYD4_CHATD/1-441
CIA1_ARATH/1-352
CIAO1_NEMVE/1-327
CIAO1_CRYNB/1-440
CIAO1_XENTR/1-333
CIAO1_DICDI/1-333
CIAO1_DANRE/1-330
CIAO1_DROME/1-335
CIAO1_NEUCR/1-468
CFD1_SCHPO/1-608
CIAO1_CAEEL/1-337
A0A1D6HWP7_MAIZE/1-333
A0A7M7GGD2_STRPU/1-340
A0A8V0ZD81_CHICK/1-342
A0A9P6YE26_RHIOR/1-342
B8CDW9_THAPS/1-414
F6VEB2_CIOIN/1-337
Q23K67_TETTS/1-342
Q581A6_TRYB2/1-424
Q6YVM7_ORYSJ/1-353
S4RIF4_PETMA/1-340

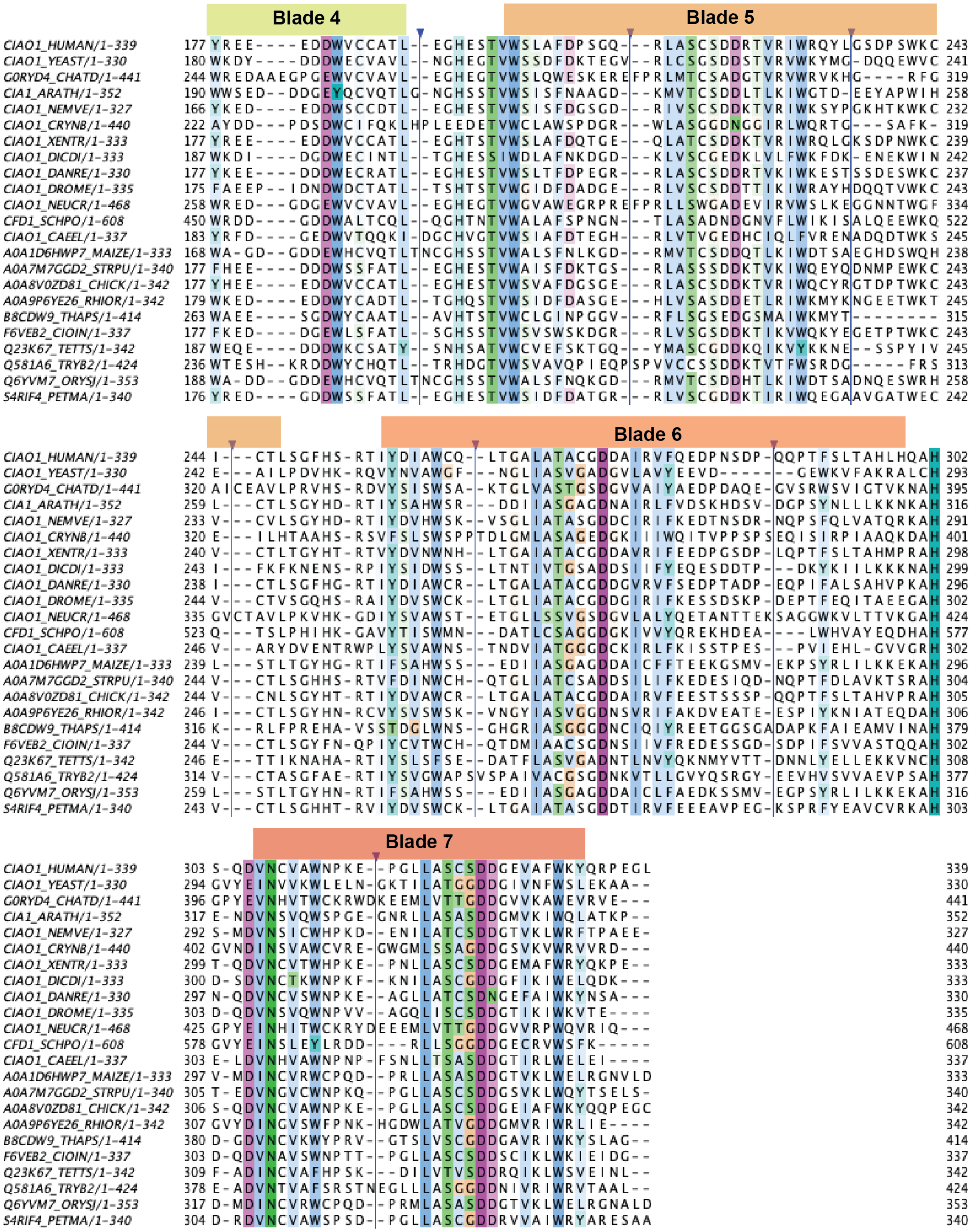


**Figure S5. Multiple sequence alignment of Cia1.** ClustalO alignment of Cia1 orthologs (Uniprot identifiers indicated) from a representative eukaryotic lineages, including: **animals** [*Homo sapiens* (HUMAN), *Nematostella vectensis* (NEMVE), *Xenopus tropicali*s (XENTR), *Danio rerio* (DANRE), *Drosophila*

*melanogaster* (DROME), *Caenorhabditis elegans* (CAEEL), *Strongylocentrotus purpuratus* (STRPU), *Gallus gallus* (CHICK), *Ciona intestinalis* (CIOIN), *Petromyzon marinus* (PETMA)], **fungi** [*S. cerevisiae (*YEAST), *C. thermophilum* (CHATD), *Cryptococcus neoformans*, (CRYNB), *Neurospora crassa* (NEUCR), *Schizosaccharomyces pombe* (SCHPO), *Rhizopus oryzae* (RHIOR)], **plants** [*Arabidopsis thaliana* (ARATH), *Zea mays* (MAIZE), *Oryza sativa* (ORYSJ)], and **other lineages** [*Dictyostelium discoideum* (DICDI), *Thalassiosira pseudonana* (THAPS), *Trypanosoma brucei* (TRYB2)]. The seven blades in β-propeller are indicated with colored rectangles. The region corresponding to **Figure 4D** is boxed in red. Blue triangles at the top indicate regions where extensions present in only a handful of orthologs were hidden for clarity. Alignments were colored with Jalview and used in Consurf server[41] to color Cia1 surface shown in **Figure 4C.**

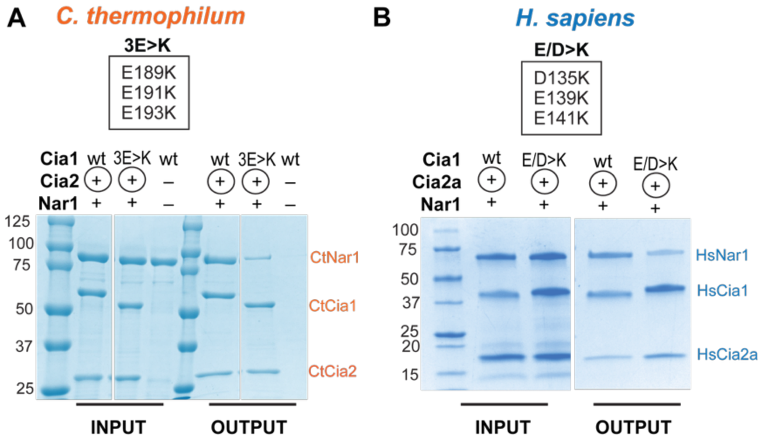


**Figure S6: Charge-reversal variants within the conserved acidic surface of Cia1 disrupt interaction with Nar1.** SDS-PAGE analysis (Coomassie staining) of affinity copurification assays monitoring the interaction of Nar1 with Cia1 variants from *C. thermophilum* (**A**) or *H. sapiens* (**B**). In all cases, Cia2 is the strep-tagged bait (circled).

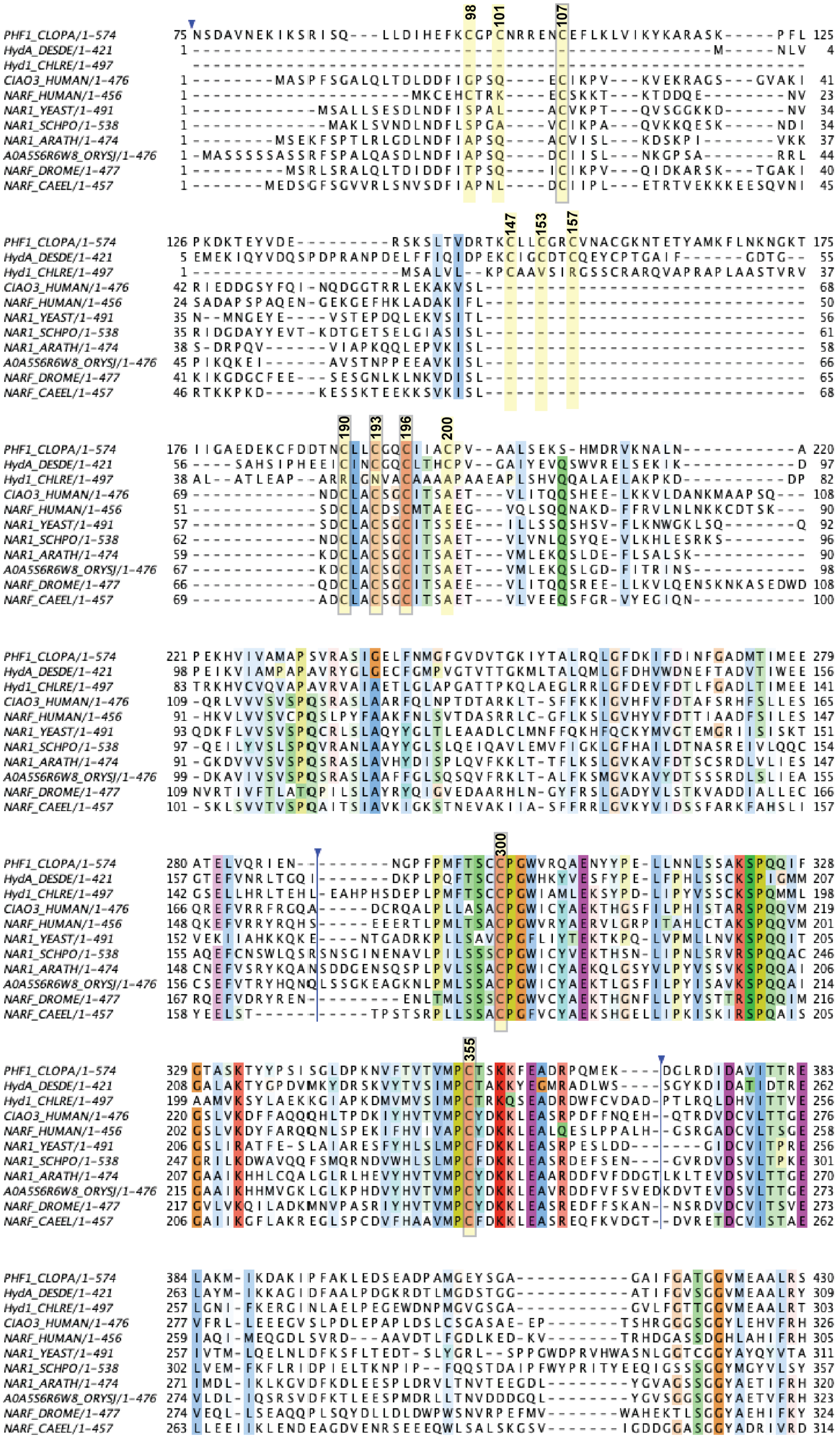

98
101
107
147
153
157
190
193
196
200
300
355
PHF1_CLOPA/1-574
HydA_DESDE/1-421
Hyd1_CHLRE/1-497
CIAO3_HUMAN/1-476
NARF_HUMAN/1-456
NAR1_YEAST/1-491
NAR1_SCHPO/1-538
NAR1_ARATH/1-474
A0A5S6R6W8_ORYSJ/1-476
NARF_DROME/1-477
NARF_CAEEL/1-457

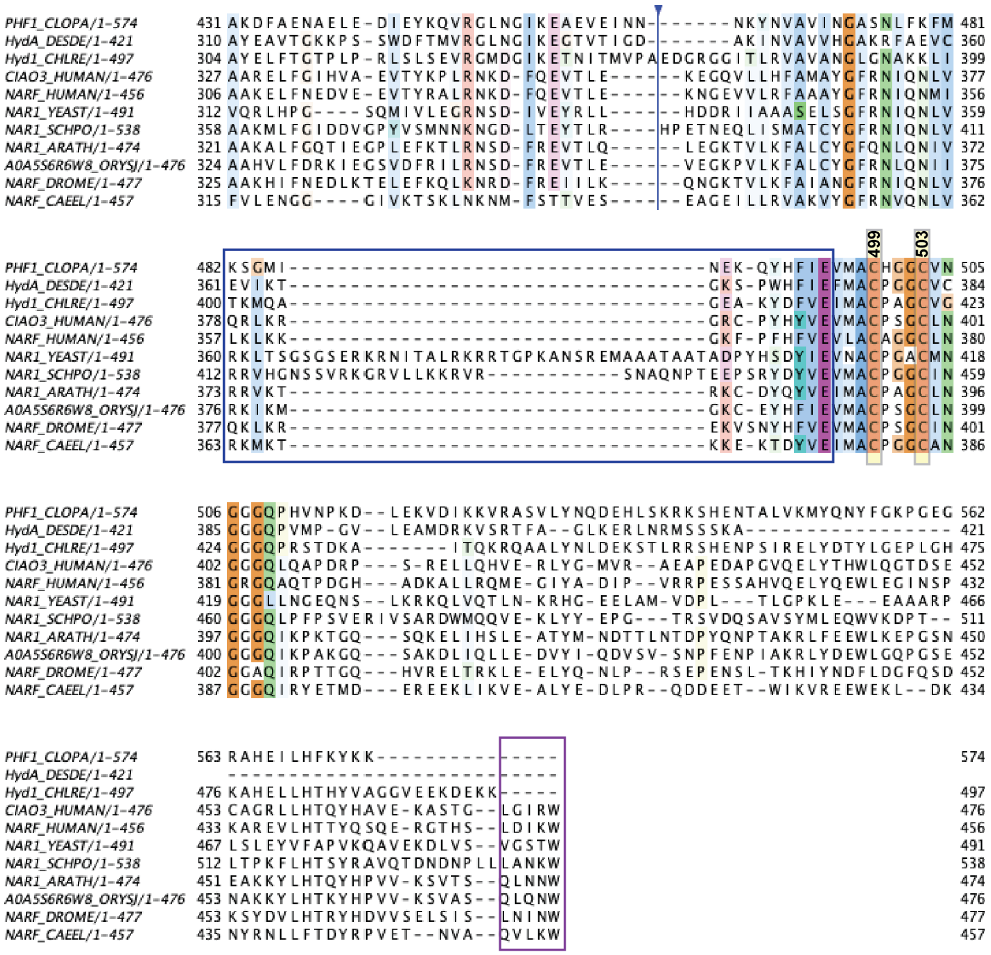

**Figure S7. Sequence alignment of [FeFe]-Hydrogenase with representative Nar1 sequences.** Hydrogenase sequences correspond to available FeFe-hydrogenase structures, PDB ID: 6N59; 1HFE; 3LX4, for PHF1_CLOPA, HYDA_DESDA, HYD1_CHLRE, respectively**.** Uniprot identifiers for the Nar1/NarF homologs derived **animals** [*Homo sapiens* (HUMAN), *Drosophila melanogaster* (DROME), *Caenorhabditis elegans* (CAEEL)], **fungi** [*S. cerevisiae (*YEAST), *Schizosaccharomyces pombe* (SCHPO)], **plants** [*Arabidopsis thaliana* (ARATH), *Oryza sativa* (ORYSJ)]. Two Nar1 paralogs from humans (CIAO3_HUMAN and NARF_HUMAN) are displayed to show homology of Nar1 proteins with the NarF paralog found in some vertebrates. Residues coordinating the [Fe–S] clusters of PHF1_CLOPA are boxed in yellow. The region enriched in positively charged residues unique to Nar1 proteins is boxed in blue. The C-terminal targeting complex recognition motif is boxed in purple.

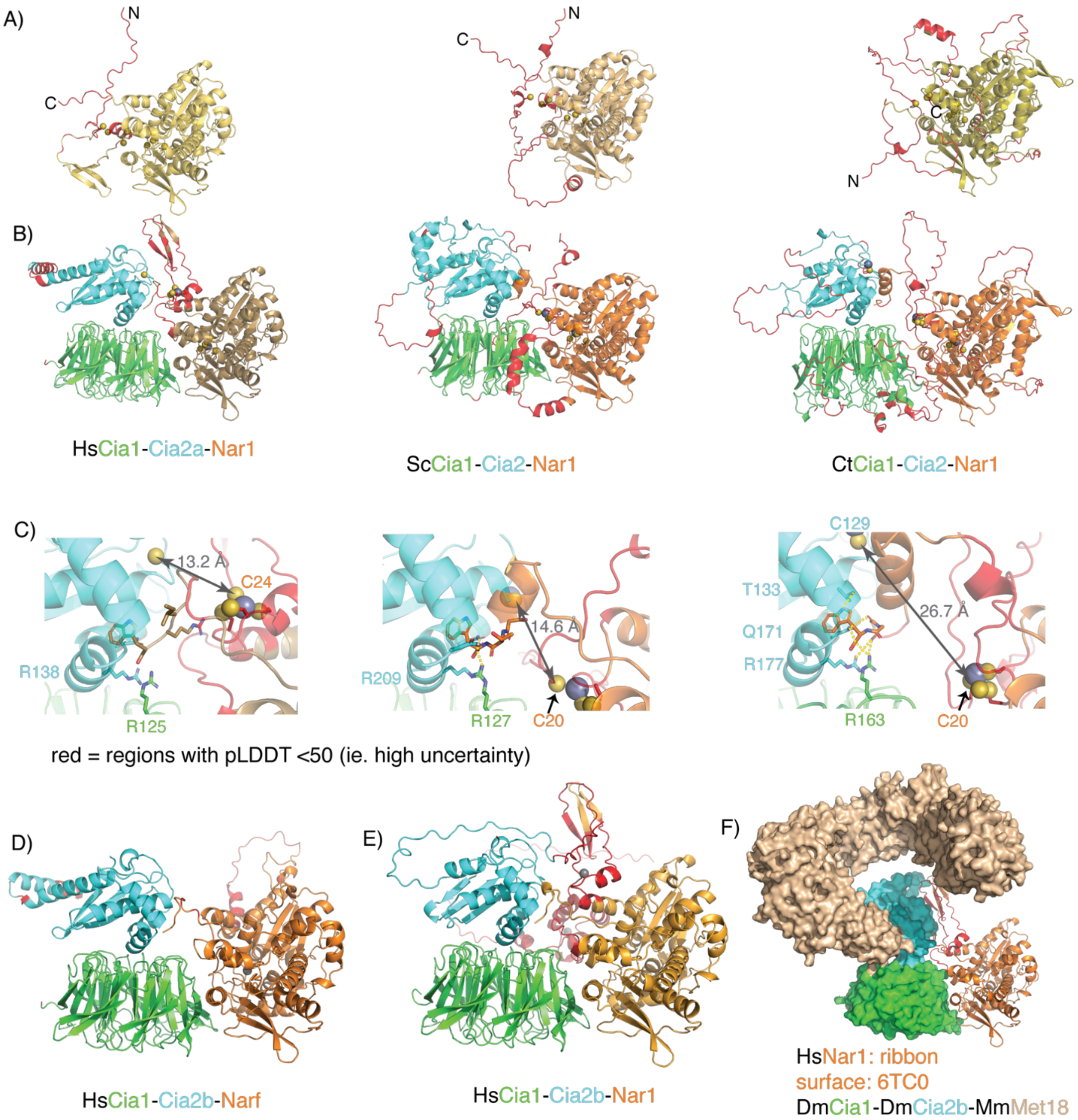


**Figure S8. AlphaFold3 modeling of Nar1 and CCN assemblies, related to Figure 5. (A)** AlphaFold3 predicted structures of Nar1 from *H. sapiens* (left), *S. cerevisiae* (middle) and *C. thermophilum* (right). Regions of high uncertainty (pLDDT <50) are red. The N- and C-termini are indicated with N and C, respectively. (**B)** AlphaFold3-models of the Cia1-Cia2-Nar1 complexes with Cia1 colored green, Cia2 in cyan and Nar1 in orange. **(C)** A close-up view of the Nar1 C-terminus (orange sticks) ound at the Cia1 (green) Cia2 (cyan) interface. The distance between the putative Nar1 Fe–S donor and the proposed Cia2 acceptor site is indicated with the double sided arrow. **(D)** HsCCN with HsNarF in place of HsNar1. **(E)** overlay of the experimentally determined CTC monomer (PDB ID 6TC0, surface) with the AlphaFold model for the HsCia-Cia2a-Nar1 complex (panel B, left). The *Hs*Nar1 is in orange ribbons to illustrate putative docking of Nar1 onto the full targeting complex given its experimentally determined structure.[17]

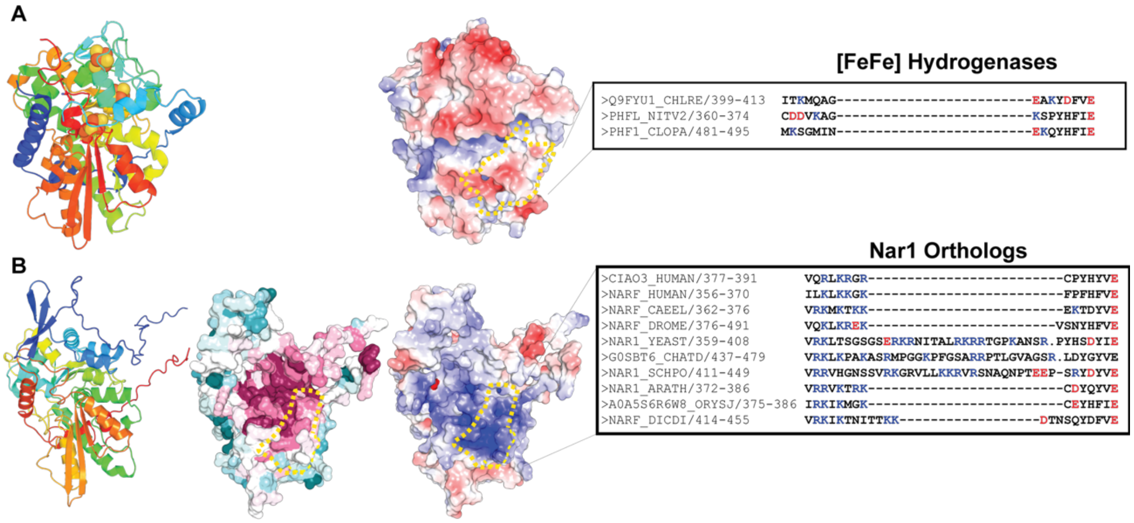


**Figure S9**. **Structural and electrostatic comparisons of [FeFe]-hydrogenase and Nar1.** (**A**) Structure of a representative [FeFe]-hydrogenase (PDB ID: 1HFE) shown as a ribbon diagram (left, colored blue to red from N- to C-terminus) with its three Fe–S clusters shown as spheres. The panel to the right shows the same protein in the same orientation colored by electrostatic surface potential. The yellow dashed line indicates the region corresponding to the basic patch identified in Nar1, together with the corresponding primary sequence region from three representative [FeFe]-hydrogenases. (**B**) AlphaFold3 model of *Hs*Nar1 shown in a similar orientation as [FeFe]-hydrogenase structure in panel **A** to highlight the structural similarities between the two proteins. The panels to the right show the *Hs*Nar1 surface colored by conservation (ConSurf, pink to cyan indicates high to low conservation)[41] and electrostatic potential.[39,40] The yellow dashed line outlines the basic surface patch and its corresponding primary structure in a representative set of Nar1 homologs.

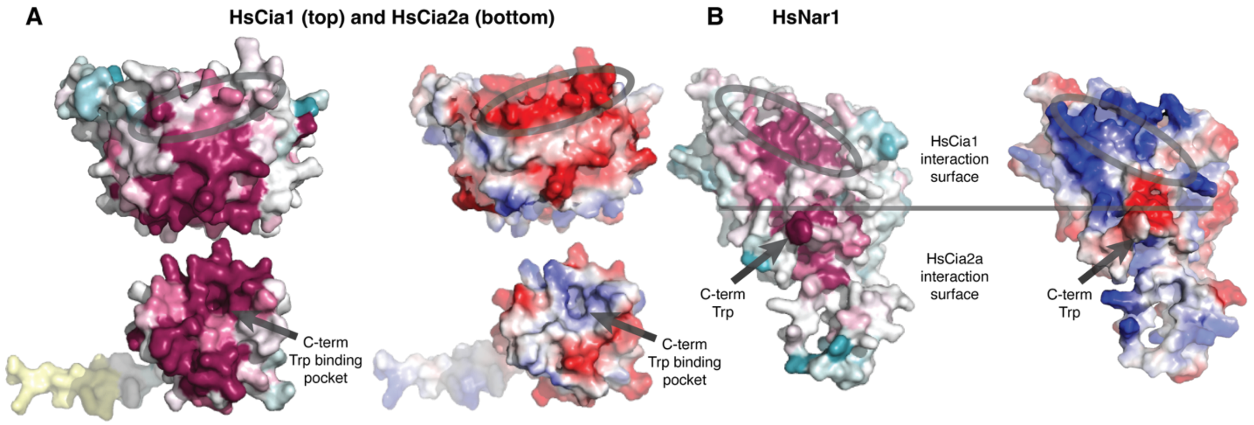


**Figure S10**. Protein-protein interaction interfaces in AlphaFold3 model of the *Hs*CCN complex. (**A**) Surface representation of *Hs*Cia1 (top) and *Hs*Cia2a (bottom) corresponding to the Nar1-interacting interface. Surfaces colored by sequence conservation (ConSurf, left) and electrostatic potential (right). (**B**) Surface representation of Nar1 surface interacting with Cia1-Cia2a colored by conservation (left) and conservation (right). The grey line represents the approximate location of the Cia1-Cia2 interface within the complex. In all models, the electrostatic surface mediating the Cia1-Nar1 interaction is circled, and an arrow indicates the TCR peptide (C-term Trp) and its binding pocket.

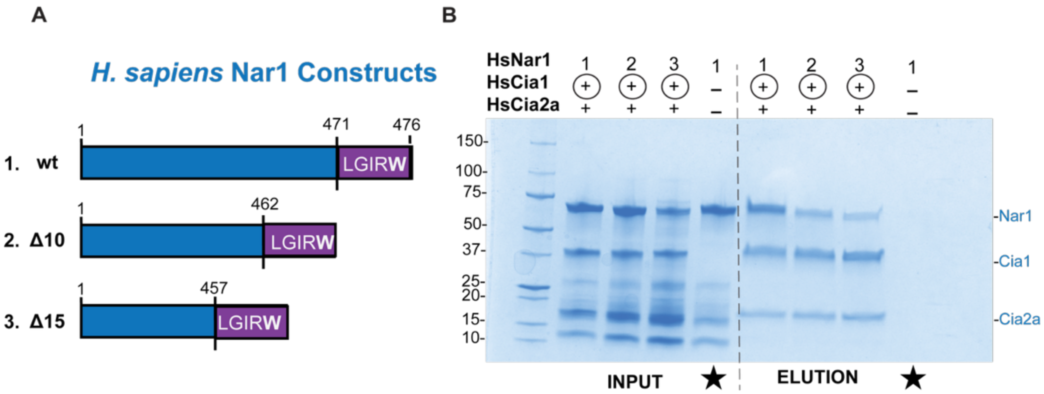


**Figure S11. Shortening of *Hs*Nar1 C-terminal tail disrupts its interaction with *Hs*Cia1-Cia2a. (A)** Schematic of the *Hs*Nar1 constructs used to probe the contribution of the C-terminal tail positioning on targeting complex binding. Constructs include the full-length protein and variants in which 10 or 15 residues were deleted (Δ10 or Δ15) between Nar1's C-terminal domain and its terminating pentapeptide (residues 471-476). (**B)** SDS-PAGE analysis (Coomassie staining) of streptactin affinity copurification assays monitoring association of the indicated *Hs*Nar1 variant with the *Hs*Cia1-Cia2a complex. In all experiments, Strep-tagged *Hs*Cia1 served as the bait (circled). Starred lanes are the no-bait control where the Strep-tagged Cia1 (and its bound Cia2a subunit) was omitted from the assay.
